# Aspen-associated soil microbiomes reveal different strategies for nitrogen acquisition across ecosystems in Mexico and Canada

**DOI:** 10.1101/2025.03.05.641730

**Authors:** Anna Fijarczyk, Roos Goessen, Marie-Josée Morency, Patrick Gagné, Jérôme Laganière, Christian Wehenkel, Javier Hernández-Velasco, Ilga Porth, Nathalie Isabel, Christine Martineau

**Affiliations:** Centre de foresterie des Laurentides, Natural Resources Canada, 1055 Rue du Peps, Québec, QC G1V 4C7, Canada; Département des sciences du bois et de la forêt, FFGG, Université Laval, 2405 Rue de la Terrasse, Québec, QC G1V 0A6, Canada; Institute for System and Integrated Biology (IBIS), Université Laval, Quebec, QC G1V 0A6, Canada; Centre for Forest Research, Université Laval, Quebec, QC G1V 0A6, Canada; Instituto de Silvicultura e Industria de la Madera, Universidad Juárez del Estado de Durango, Km 5.5 Carretera Mazatlán, 34120 Durango, Mexico; Universidad Intercultural de Baja California (UIBC), San Quintín, Baja California, C.P. 22930, México

**Author notes:** Corresponding authors: Anna Fijarczyk, Christine Martineau.

**Keywords:** *Populus tremuloides*, soil microbiome, mycorrhizal fungi, metabarcoding, nitrogen acquisition, microbial diversity

## Abstract

Plant species shape soil microbiome composition through species-specific interactions. However, it is less clear how these interactions vary across populations that diverged a long time ago. In this study, we explore the influence of host genetic composition and edaphic factors on the soil microbiome of *Populus tremuloides*, one of North America’s most widespread tree species. Using 16S, 18S rRNA gene, and ITS2 region metabarcoding on soils from natural stands and potting mix, rhizosphere, and root samples from a greenhouse common garden, we examined prokaryotic and fungal communities in two aspen genetic groups. The Eastern Canada group represents boreal and cold temperate ecoregions, and the one from Northwestern Mexico represents warm temperate ecoregion. Variation in microbial community structure correlated with soil properties but results from common gardens indicated that the host genetic makeup may also play a role. The three ecoregions showed functional divergence: warm temperate sites hosted a higher abundance and diversity of nitrogen-fixing bacteria, while boreal stands exhibited stronger associations with ectomycorrhizal fungi. Our findings highlight how local adaptations to climate and soil conditions in aspen extend to their microbial partners, emphasizing the potential role of host-microbe interactions in shaping tree resilience and susceptibility to future climate changes.

## Introduction

Forests play a critical role in carbon cycling, particularly as carbon sinks, with their microbiota being a key component in this process (Malhi, Baldocchi and Jarvis 1999; Baldrian 2017).

Microorganisms act as carbon storage, they decompose organic material and provide vital services to forest species by supplying nitrogen and phosphorus or promoting tree growth (Uroz *et al*. 2016; Baldrian 2017; Hawkins *et al*. 2023). At the same time, forest microbiomes also harbor pathogens that affect species population dynamics and forest services (Castello, Leopold and Smallidge 1995; Boyd *et al*. 2013). The diversity and dynamics of microbial communities are key to understanding the functioning of forest ecosystems.

Globally, climate, growing season length, and edaphic factors shape forest species composition and their microbial communities (Baldrian, López-Mondéjar and Kohout 2023). Along latitudinal gradients, forest ecosystems vary in species richness, macronutrient availability, and organic matter content (Malhi, Baldocchi and Jarvis 1999; Reich *et al*. 2014; Gao *et al*. 2022; Liang *et al*. 2022). Boreal forests are generally low in eukaryotic species diversity and nutrients but have abundant organic matter (Pan *et al*. 2013; Liang *et al*. 2022). In contrast, tropical forests are rich in species and nutrients but contain less organic matter (Pan *et al*. 2013; Liang *et al*. 2022).

Nutrient availability and tree species composition have a strong impact on the composition of microbial communities in the soil (Uroz *et al*. 2016; Baldrian, López-Mondéjar and Kohout 2023). These include symbiotic components of the soil microbiome, such as ectomycorrhizal fungi (EMF) and arbuscular mycorrhizal fungi (AMF), that are responsible for the transport of nitrogen and phosphorus to plants in exchange for carbon via root connections (Courty *et al*. 2010; Powell and Rillig 2018; Ward *et al*. 2022). Mycorrhizal fungi exhibit diverse host preferences; they influence both forest growth and biodiversity, making them crucial to the overall health of the forest.

*Populus tremuloides* Michx. is one of the few species in North America with a distribution range that spans multiple ecosystems and forms both ectomycorrhizal and arbuscular mycorrhizal associations (Teste, Jones and Dickie 2020). *P. tremuloides* populations extend across diverse ecological regions: they dominate much of the boreal zone in the north, stretching into the eastern temperate zone in the middle-eastern part of North America, whereas in the west they spread through the high-elevation northwestern forested mountains, reaching as far south as the temperate Sierras of the United States of America and Mexico. Aspens are keystone forest species supporting wildlife habitats and contributing to ecosystem diversity (Rogers *et al*. 2020), but a decline in populations has been observed in recent years (Worrall *et al*. 2013), with drought identified as a key factor in the reduced growth of Western Canadian populations (Chen *et al*. 2017).

Genetically, *P. tremuloides* is divided into four major genetic groups spread across distinct regions: Eastern Canada, Western Canada, the Midwestern and Southern U.S.A., and Mexico, each varying in levels of clonality and ploidy (Goessen *et al*. 2022), although more groups can be distinguished with more extensive sampling (Goessen, unpublished). In the southwestern range (U.S.A. and Mexico), large clonal stands are common (Kemperman and Barnes 1976; Mitton and Grant 1996), with the Pando clonal population in the Fishlake National Forest (U.S.A.) being an extreme example (DeWoody *et al*. 2008; Pineau *et al*. 2024). In areas like the Great Lakes and Central Canada, sexual reproduction occurs more frequently, contributing to smaller clone sizes (Namroud *et al*. 2005; Mock *et al*. 2008), and seed germination being less affected by drought (Goessen *et al*. 2022). Given their broad ecological distribution, the varying vegetation composition and density across regions, local climate adaptations within genetic groups are expected (Goessen *et al*. 2022). The range of naturally occurring microbial associations across these groups is still unknown. In a large-scale study comprising several aspen species, climate and geography have been shown to be the primary drivers of soil and foliar endophytic fungal communities, with arbuscular mycorrhizae more prevalent in warmer regions and ectomycorrhizal fungi more common in cooler areas (Van Nuland *et al*. 2023).

However, to the best of our knowledge, no study has yet compared the microbiomes within aspen stands belonging to a single species across large-scale ecosystems, such as boreal and temperate zones.

Here we use amplicon sequencing of 16S rRNA gene, ITS2 region, and 18S rRNA gene to investigate soil prokaryotic, fungal, and arbuscular mycorrhizal fungal microbiomes, respectively, in natural stands (n=46) and greenhouse settings (n=12) of *P. tremuloides* coming from two distinct genetic groups: Eastern Canada and Northwestern Mexico. Knowing that the two genetic groups of aspen show phenotypic evidence for local adaptation (Goessen 2023), we hypothesise that relative abundances and diversity of soil microbial communities including symbiotic microorganisms associated with the two genetic groups of aspen have differentiated. The two genetic groups span three ecoregions: populations from Canada span boreal and cold temperate zones, whereas populations from Mexico, are located in the warm temperate zone.

Because global patterns of symbioses differ across ecoregions (Steidinger *et al*. 2019), we also expect soil communities to vary between the three regions. Our research objectives were to: i) examine the differences in microbial communities between aspen-dominated stands in Canada and in Mexico and determine how soil and aspen genotypes influence them, ii) assess the abundance and diversity of EMF, AMF and nitrogen-fixing bacteria in these groups, and iii) evaluate the effect of aspen’s genetics on microbial community structure, including EMF, AMF communities and nitrogen-fixing bacteria, in a greenhouse common garden setup.

## Materials and Methods

### Study sites

Forty-six soil samples were collected at eight sites within a two-meter radius of *P. tremuloides* trees growing in patchy stands, generally within mixed forests (Fig. 1A-C, Supplementary Tables 1-3). Sampling was conducted at five sites in Quebec, Canada, and three sites in Northwestern Mexico (Fig. 1B). In Canada, two sites (AMOS, STFE) are located well within the boreal zone, while one site (STET) falls within the Mixedwood Plains ecozone corresponding to the cold temperate region. The remaining two sites (ESSI, FORE) lie at the southern limit of the boreal zone, in the Saint Lawrence Estuary, where soil and climate characteristics resemble those of a temperate region (Fig. 1E). Given these similarities, we grouped STET, ESSI, and FORE into a cold temperate region (Fig. 1E). Sampling in Northwestern Mexico was done at three sites within the warm temperate zone (Santiago, FP2, and FLOR1, Fig. 1, Supplementary Figure 1). One additional site, AB within the boreal zone in Canada, was included in the greenhouse experiment only (Fig. 1A-B,D). Sites were accessed either in 2018 (ESSI, AMOS, STFE, FORE), or in 2019 (the rest of sites).

**Fig 1.**
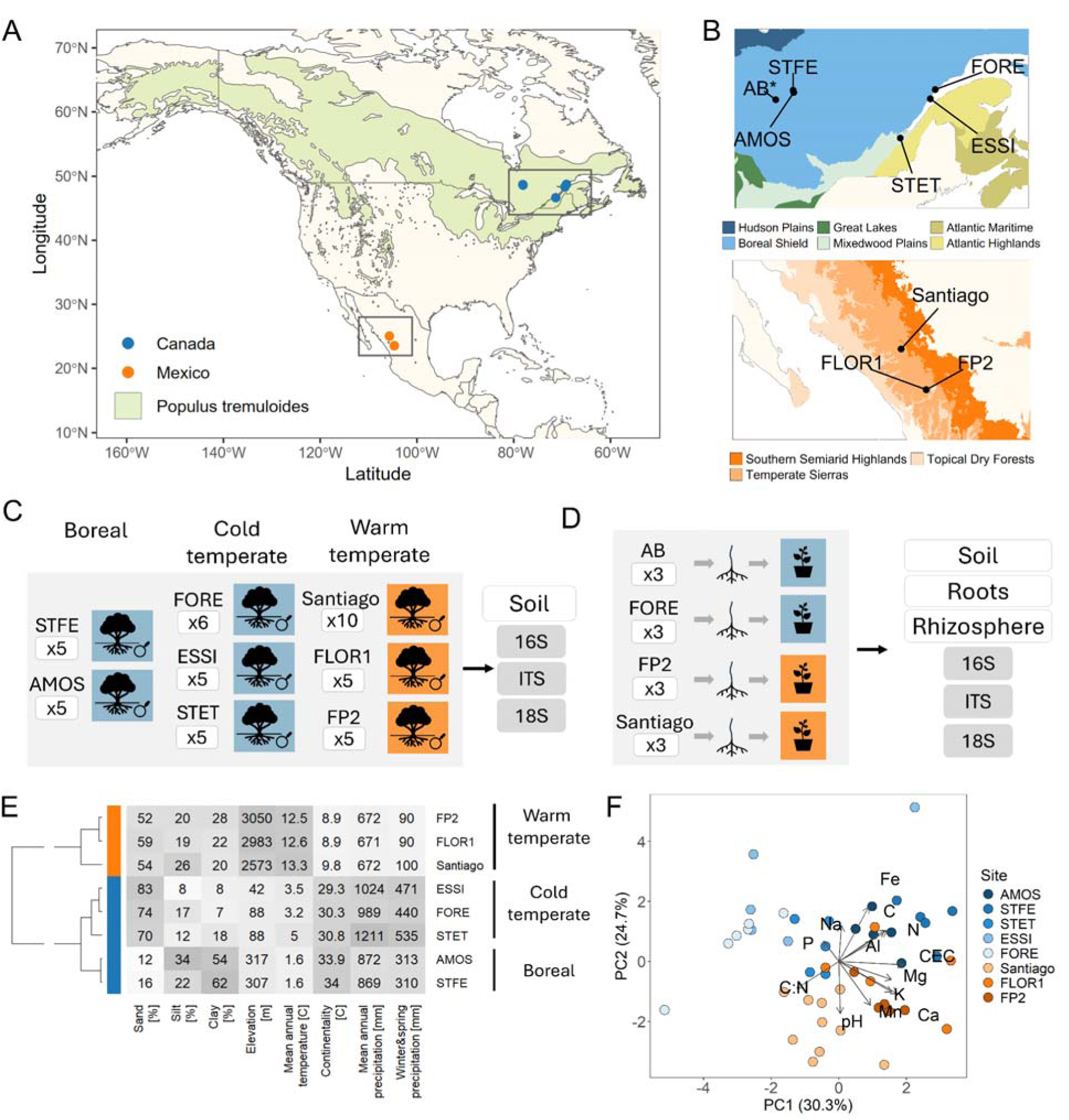
Study design and characteristics of the sampling sites. A) Distribution of aspen with the sampling locations. B) Sampling locations in Quebec, Canada and Northwestern Mexico. Note that site AB was used for the greenhouse study only. C) Datasets collected from the natural stands. D) Design of the greenhouse setup and analyzed datasets. E) Soil texture and geoclimate characteristics in seven natural stand locations. Climate variables represent 1990–2020 climate normals. F) Principal component analysis of soil physicochemical properties. Ecoregions displayed in panel (B) were obtained from the Government of Canada website (https://natural-resources.canada.ca/forest-forestry/sustainable-forest-management/forest-classification) and the United States Environmental Protection Agency website (https://www.epa.gov/eco-research/ecoregions-north-america). Distributions and tests of all soil properties are provided in Supplementary Figure S2.

The sites AMOS, STFE, and AB are part of the Canadian clay belt in Quebec, Canada. Apart from soils rich in clay and silt, these sites are characterized by the lowest mean annual temperatures (1.6॰C) among all sampled sites, and relatively high mean annual precipitation (869-872 mm). Sites AMOS (Guillemette and DesRochers 2008) and STFE are situated in aspen natural stands, with understory vegetation including plants such as ferns, *Rubus* spp., *Epilobium* spp., and seedlings of *Salix* spp., *Betula* spp., and *Abies balsamea*. The AB site is part of a natural post-fire mixedwood forest, featuring species such as *A. balsamea*, *Picea mariana*, *P. glauca*, and *B. papyrifera* (Bergeron 2000).

Three Canadian sites ESSI, FORE, and STET are located along the Saint Lawrence Estuary. These sites are located on sandy soils, at low elevations, and experience high mean annual precipitation of up to 1211 mm. ESSI and FORE sites consist of natural mixed stands of *P. tremuloides* and conifers, situated near the roadside with varying levels of disturbance. The STET site is characterized by natural stands of *P. tremuloides* growing on abandoned farmland near a river and residential areas.

The three Mexican sites—Santiago, FP2, and FLOR1—are all located in Durango State in the Sierra Madre Occidental range (temperate Sierras zone), within pine-oak forests. The sites are found at high elevations (2573-3050 m) with annual mean temperatures around 12.5-13.3॰C and mean annual precipitation reaching 672 mm. In contrast to Canadian sites, these locations experience less temperature fluctuations throughout the year, whereas precipitation is seasonal.

Rainfall occurs mostly in summer and autumn with long periods of dry weather in between. The FLOR1 and FP2 sites are dominated by *Arbutus bicolor*, *Arbutus* spp., *Pinus durangensis*, *Pinus arizonica*, and *Quercus rugosa*. The Santiago site is situated on soils with sparse vegetation near a road, and a stream present only during the rainy season. Its natural stands are dominated by *Pinus arizonica*, *Quercus sideroxyla*, and *Arbutus* spp.

### Sampling protocol

Soil was taken in triplicates at 2 m distance from the tree. Samples were obtained using a 15 cm soil corer after removing the organic layer and stored in plastic bags kept on ice in a cooler.

Sampling equipment was cleaned to remove residual soil and wiped with ethanol between each sample. In the lab, triplicates were mixed into a composite sample and further homogenized using a 6 mm mesh sieve. Root segments were removed from the soil samples. A 0.25 g subsample of the resulting soil was transferred to a PowerBead tube (DNeasy PowerSoil Kit, QIAGEN) for DNA extraction. The remaining soil was air-dried and used for physicochemical analyses.

### Greenhouse experiments

*P. tremuloides* saplings were grown in a greenhouse compartment at Laval University’s Forestry faculty in Quebec City, from the roots collected at two locations in Quebec (AB, FORE) and two locations in Mexico (FP2 and Santiago). Samples from Quebec were received in September 2019 and were first propagated by putting them on trays with peat moss soil to stimulate root sprouting and then transferring cuttings of around 10 cm to a misting unit on perlite substrate for rooting. After developing roots, plants were transferred to the potting mix. Roots from Mexico were received 1-2 months after harvest (November 2019 and January 2020 for FP2 and Santiago, respectively). Due to the low propagation success rate of previous batches of Mexican roots, these roots were planted directly in a potting mix. The substrate for growing cuttings included 2 volumes of peat (Fafard), 1 volume of perlite, and 0.5 volume of vermiculite.

Cuttings were repotted several times and transferred to 1.7 L pots between May and June 2020, and to 2.8 L pots between January and February 2021. The substrate for repotting was composed of 3 volumes of peat (Fafard), 1 volume of perlite, 1 volume of vermiculite, and 0.25 volume of compost (Biosol, de Fafard). Several pesticide treatments were used including Nova 40 WP (September 2020, August 2021), Bioceres (September 2020), and Avid (January 2021), and predator insects (Cucumeris, Dalotia, Orius, and Aphidus) were used to limit pests. Plants were fertilized weekly with nitrogen, phosphorus, and potassium (20-20-20, 1 gL^-1^) and micronutrients (Plant-Prod Chelated Micronutrient Fertilizer Mix, 0.3 gL^-1^). Overall, plants were grown under ambient temperature and long daylight in peat moss soil for 21-24 months.

Potting mix, root, and rhizosphere samples were collected from pots on 12 October 2021 and subsequently transferred to and processed in the research laboratory at the Laurentian Forestry Centre (Canadian Forest Service, Quebec City) to obtain DNA extracts. Potting mix was processed as soil above, whereas root and rhizosphere samples were obtained for DNA extraction using the following protocol. Roots with attached substrate (rhizosphere) were transferred to a 25 mL solution containing 1X PBS (phosphate-buffered saline) and Tween 20 (0.1%) and shaken by inversion 10 times to detach particles from the roots. Tubes were then centrifuged at 4500 g for 10 minutes at 4°C. Roots were transferred to a petri dish half-filled with a fresh PBS solution and rinsed by pulling them gently through the solution with sterile tweezers. Clean roots were transferred to a 50 ml Falcon tube containing 25 ml PBS, submerged, vortexed for 15 seconds, and put twice in a sonication bath. Each sonication step was done at room temperature and at an operating frequency of about 40 kHz for 30 seconds 5 times with 30-second intervals between repetitions. Before the second sonication, roots were transferred to clean tubes containing 25 ml PBS. Roots were recovered from the tubes, pat-dried on absorbent paper and cut into 1 cm pieces with scissors pre-sterilized in 70% ethanol.

Subsequently, 0.1 g of the root fragments were transferred to a 2 ml tube with a 0.5 mm steel bead and stored in the freezer (at-20°C). Frozen roots were homogenized twice for 45 seconds each on the TissueLyser II (QIAGEN) and collected for DNA extraction. The rhizosphere pellet was recovered from the tube after centrifugation of the roots with attached substrate, after removing roots and the supernatant. The pellet was thoroughly mixed with a sterilized spatula and transferred to a small piece of Whatman filter paper to absorb the remaining PBS liquid.

Finally, 0.25 g of rhizosphere was taken for DNA extraction.

### DNA extraction and library preparation and sequencing

DNA extraction from soil, potting mix, rhizosphere, and root samples was performed using the DNeasy PowerSoil Kit (QIAGEN) following the manufacturer’s instructions. DNA concentration of the extracts was assessed with the Qubit 3.0 fluorometer (Thermo Fisher Scientific, MA, USA) using the dsDNA BR (broad range) or HS (high sensitivity) assay kit (Life Technologies, CA, USA). Amplicon library preparation was performed as described by Illumina (Illumina 2013) using user-defined primers. Selected primer pairs were 515 F-Y (5′-GTGYCAGCMGCCGCGGTAA-3′) and 926R (5′-CCGYCAATTYMTTTRAGTTT-3′) targeting the V4–V5 regions of the 16S rRNA gene of bacteria and Archaea (Parada, Needham and Fuhrman 2016), and ITS9F (5′-GAACGCAGCRAAIIGYGA-3′) and ITS4R (5′-TCCTCCGCTTATTGATATGC-3′) targeting the ITS2 region of fungi (White *et al*. 1990; Menkis *et al*. 2012). PCR reactions were set up by mixing 25.0 µL of 2X HotStarTaq Plus Master Mix (QIAGEN, Germany), 0.5 µL of each 10 µM HPLC-purified (or desalted) primer (Invitrogen, USA), 19 µL of RNase-free water (QIAGEN) and 5.0 µL of DNA diluted at 5 ng µL^-1^, for a total volume of 50 µL. Thermal cycling conditions for the 16S rRNA gene and ITS2 region were as follows: initial denaturation at 95°C for 5 min; 35 cycles or 40 cycles for ITS2 at 94°C for 45 s, 50°C for 45 s, 72°C for 1 min; and a final elongation step at 72°C for 10 min. For the 18S rRNA gene, targeting AMF, a nested PCR approach was used, as previously described (Stefani *et al*. 2020). In the first step, samples were amplified using primers AML1 (5′-ATCAACTTTCGATGGTAGGATAGA-3′) and AML2 (5′-GAACCCAAACACTTTGGTTTCC-3′) targeting the V3-V4-V5 region of the 18S rRNA gene with the following thermal cycling conditions: 95°C for 5 min; 35 cycles of 94°C for 1 min, 57°C for 1 min, 72°C for 1 min; and a final elongation step at 72°C for 10 min. PCR products were used as template in a second PCR reactions with primers nu-SSU-0595-5’ (5′-CGGTAATTCCAGCTCCAATAG-3′) and nu-SSU-0948-3′ (5′-TTGATTAATGAAAACATCCTTGGC-3′) targeting the V4 region of the 18S rRNA gene with the following thermal cycling conditions: 95°C for 5 min; 35 cycles of 94°C for 40 sec, 57°C for 40 sec, 72°C for 40 sec. Amplicon libraries were sequenced on an Illumina MiSeq platform using a PE300 v3 kit at the Centre de recherche du CHU de Québec-Université Laval. Samples were sequenced in three runs: i) 16S rRNA gene and ITS2 region of all samples from natural stands, ii) 16S rRNA gene and ITS2 region of all samples from the greenhouse experiment, iii) 18S rRNA gene of all samples.

## Data acquisition and processing

### Climate and soil physicochemical properties

For each sampling site, we used ClimateNA v7.50 (Wang *et al*. 2016) to obtain annual mean temperature, annual mean precipitation, continentality (the difference between the mean of the warmest month and mean of the coldest month), and mean precipitation for winter and spring seasons, for years between 1991 and 2020. Elevation was extracted for each site using Python package *pyhigh* v0.0.6 (https://github.com/sgherbst/pyhigh/).

Soil obtained through sieving with 6 mm mesh was air-dried and subjected to physicochemical analyses. Soil was further sieved with 2 mm mesh size, and a soil subsample was additionally ground to obtain a 0.5 mm fraction using a disk mill for the determination of percent total carbon (C) and nitrogen (N). N and C were quantified with TruMac CNS analyzer (LECO Corporation, MI, United States). Using a 1:2 soil:water ratio, the soil pH was determined in the aqueous phase with a Thermo Scientific Orion 2-Star pH meter following (Carter and Gregorich 2007). Phosphorus (P), potassium (K), calcium (Ca), magnesium (Mg), manganese (Mn), iron (Fe), aluminum (Al), and sodium (Na) were extracted with a Mehlich III extraction buffer (Carter and Gregorich 2007) and quantified using an ICP-OES (Optima 7300 DV, Perkin Elmer, Waltham, MA, United States). Cation exchange capacity (CEC) was calculated using quantified positively charged ions. The soil texture (sand, silt, and clay) was determined following the hydrometer method (Carter and Gregorich 2007).

### Bioinformatic analysis

Reads (16S, 18S rRNA genes, and ITS2 region) were processed using the Q2Pipe pipeline (https://github.com/NRCan/Q2Pipe, versions 10-2022 (natural stands), 01-2024 (greenhouse samples), and 05-2024 (18S rRNA gene)), which leverages the functionality of the QIIME2 platform (Bolyen *et al*. 2019). In short, read quality was initially evaluated using Figaro (Weinstein *et al*. 2019) and Falco (https://zenodo.org/badge/latestdoi/214499063) to identify trimming positions to remove adapters and low-quality positions for 16S and 18S rRNA gene targets. After trimming, the 16S and 18S rRNA gene reads were filtered, and merged into Amplicon Sequence Variants (ASVs) with DADA2 (Callahan *et al*. 2016). Because the length of the ITS2 region can vary, instead of positional trimming, ITS2 region reads were trimmed for adapters using CutAdapt (Martin 2011) and then filtered and merged into ASV as described above. ASVs with frequencies below 0.05% were discarded, and the remaining ASVs were assigned to taxonomic levels. For taxonomic classification, we used databases UNITE v9 for the ITS2 region in fungi (Nilsson *et al*. 2019), Silva v138 for the 16S and 18S rRNA genes (Yilmaz *et al*. 2014), and FUNGuild (December 2022) for the assignment of fungal guilds (Nguyen *et al*. 2016). For the 16S rRNA genes, only sequences assigned to bacteria and Archaea were retained, excluding sequences classified as Eukaryota, mitochondria, or chloroplast. For the ITS2 region, we retained sequences classified as fungi, and for the 18S rRNA gene those classified as Eukaryota, since the 18S rRNA gene primers target AM fungi but are not 100% specific. In natural stands we retained a total of 1,423,526 reads (16S rRNA gene), 1,555,740 reads (ITS2 region), and 1,677,810 reads (18S rRNA gene) for further analyses. For the greenhouse dataset, we retained 739,825 reads (16S rRNA gene), 2,082,460 reads (ITS2 region), and 1,671,470 reads (18S rRNA gene). For estimating alpha diversity, read counts were rarefied using rarefaction curves as a guide to control for uneven read numbers across samples. Read counts of natural stand samples were rarefied to 6370, 11982 and 19500 for the 16S rRNA gene, ITS2 region, and 18S rRNA gene datasets, respectively. Read counts of greenhouse samples were rarefied to 7921, 12594 and 19500 for 16S rRNA gene, ITS2 region, and 18S rRNA gene datasets, respectively. Other analyses were conducted on unrarefied datasets using statistical approaches dedicated to the analysis of compositional data, including transformation of read counts, all of which are described in detail below.

## Statistical analyses

All downstream analyses were performed with R v4.4.1 (R Core Team 2021) and Python v3.8 (Van Rossum and Drake 2009).

In the natural stands dataset, four samples were excluded from the ITS2 analysis due to low read counts (1x STET and 3x Santiago), resulting in 46 samples with 16S rRNA gene sequences, 42 samples with ITS2 region sequences, and 46 with 18S rRNA gene sequences. The greenhouse dataset included 12 potting mix, 12 rhizosphere, and 12 root samples, all of which were sequenced for 16S, 18S rRNA genes and ITS2 regions. Nitrogen fixing bacteria were identified based on taxonomic classification, by searching for both symbiotic and free-living bacterial genera that are known to be capable of nitrogen fixation. We included all ASVs for which the genus name contained the word “rhizobium” or “Rhizobium”, all ASVs from the phylum Cyanobacteria, and genera such as *Frankia*, *Ensifer*, *Azotobacter*, *Azospirillum*, *Beijerinckia*, or *Azoarcus* Rhizobia, *Frankia* and Cyanobacteria were the most commonly found taxa.

Differences in each soil property between genetic groups, ecological regions and sampling sites were tested with one-way ANOVA with nested groups (genetic group/region/site). Tests were conducted using R function anova() after fitting linear models with R functions lm() (package stats v.4.4.2 (R Core Team 2021)) or gls() (package *nlme* v3.1 (Pinheiro *et al*. 2025)). Outliers were removed and data was transformed when necessary, assuring normal distribution of residuals and homogeneity of variance. If conditions were not met, a generalized least square (GLS) model was used with the gls() function to account for unequal variance between the groups and correlated errors within the nested structure. GLS models were fitted separately for the site, region, and genetic group level. Benjamini-Hochberg multiple testing correction was applied to *P*-values at a false discovery rate of 0.05. Principal component analysis (PCA) on micro and macronutrients was done with R function prcomp() from the base R package stats v4.4.2 (R Core Team 2021).

Abundance tables for each taxonomic marker (16S rRNA gene, 18S rRNA gene, ITS2 region) were imported into a phyloseq R format. To visualize community composition by ordination, we retained ASVs present in 5% or more of the samples, to remove potential sequencing artifacts and reduce the noise (Cao *et al*. 2020). Read counts were then transformed to relative abundance. Principal Coordinates Analysis (PCoA) was performed on the Bray-Curtis distance matrix using the ordinate() function from the package *phyloseq* v1.50 (McMurdie and Holmes 2013). To project soil nutrient variables onto the ordination space, we conducted an envfit analysis with the R package *vegan* v2.6 (Oksanen *et al*. 2025), testing significance with 999 permutations. In the natural stands dataset, we tested the effects of grouping (genetic groups, regions, and sites) on community structure using PERMANOVA via the adonis2() function in the *vegan* v2.6 package (Oksanen *et al*. 2025), with significance tested using 999 permutations.

Permutations at the lower-level groupings (region and site) were constrained to the higher-level groupings (genetic cluster and region, respectively) using strata option. In the case of the greenhouse datasets, we tested the effects of sample type, genetic cluster (within sample type), and site (within sample type and genetic cluster) for the whole dataset and the effect of genetic cluster and site (within the genetic cluster) for each sample type using the same method.

Benjamini-Hochberg multiple testing correction was applied to *P*-values obtained for all the greenhouse datasets at a false discovery rate of 0.05.

Alpha diversity metrics were estimated from the rarefied data using the alpha() function from the *microbiome* v1.26 package (Lahti and Shetty 2012). Metrics included Gini-Simpson index (the probability that two randomly selected ASVs are different) and the total species richness (Chao1). Differences between regions were tested with one-way nested ANOVA or GLS models as described above. Pairwise differences were tested with the Tukey test. To assess the relationship between soil properties and alpha diversity (Gini-Simpson index), we fitted a beta regression model using the betareg() function from the R package *betareg* v3.2 (Cribari-Neto and Zeileis 2010). To select explanatory variables, we first selected a small subset of soil variables that were weakly correlated with each other (|r| < 0.4), but retaining those that are known to play an important role in shaping microbial diversity, and these were: pH, C:N, P, K and Na. We removed two extreme outlier points, which clearly affected correlations between variables (P > 300 mg/kg, and C:N < 70). In order to further reduce the number of explanatory variables tested in the model, we first fitted beta regression to each soil variable separately and ranked them by goodness of fit using AIC criterion. In the case of the 16S rRNA dataset, we dropped P, because it showed the lowest fit, and in the case of ITS2 region, we dropped C:N. The remaining four variables were used together with the genetic group (Mexico or Canada) as explanatory variables. We used a genetic group as a fixed effect to control for multiple factors that vary mostly between the groups and which include but are not limited to: climate variables such as temperature or precipitation, genetic ancestry of aspen, or sampling year. The best model was selected using stepwise model selection with R package *StepBeta* v2.1.0 (Garofalo 2022), starting with all possible interactions among variables. We examined diagnostics of the best models, including distribution of residuals against fitted values and collinearity. Pseudo *R*^2^ of each term in the best model was calculated by subtracting pseudo *R*^2^ of the model without the term from the pseudo *R*^2^ of the full model. In the case of the ITS2 region, the best model showed generally a poor fit (pseudo *R*^2^ = 0.12), with P and P: group as identified effects. Exploration of the model showed that it was mainly driven by P outliers, and their removal eliminated the effect of P and reduced model fit even further. We therefore concluded that a given set of soil variables cannot explain well variation in fungal diversity.

We measured the effect of the genetic distance among aspen trees on the microbiome community structure using single nucleotide polymorphism (SNP) data from (Goessen *et al*. 2022). Vcftools v0.1.16 (Danecek *et al*. 2011) was used to extract SNPs with mean depth across all individuals of more than 2, less than 15% of missing data, and a frequency of minor alleles above 5%, giving a total of 53,714 SNPs. Euclidean distances between genotypes were estimated with the dist() function from the *stats* v.4.4.2 package (R Core Team 2021), whereas for the microbial communities we calculated Bray-Curtis dissimilarities using vegdist() function in the *vegan* v2.6 package (Oksanen *et al*. 2025). To evaluate the association between genetic and microbial distance matrices we applied a Mantel test with 10,000 permutations using the mantel() function from the R package *vegan* v2.6 (Oksanen *et al*. 2025). PCA on SNPs was performed with snpgdsPCA() function from R package *SNPRelate* v1.40 (Zheng *et al*. 2012).

Differential abundance between taxa from the three sampled ecological regions (boreal, cold and warm temperate) was estimated with ANCOM-BC v2.6 (Lin and Peddada 2020), which models an unsampled fraction of taxa in compositional data. Taxa present in at least 35% of the samples were selected to remove those that occur infrequently. Differences between regions were estimated independently at the genus and family level, by first combining read counts of ASVs from the same genus or family. Taxa that were not defined or uncultured were grouped at the lowest possible taxonomic level. The sampling site was included as a random effect in the model. To assess differential abundance between regions and sites in fungal guilds and functional groups, combined read counts for the selected guilds were CLR-transformed. In the case of the 18S rRNA gene, we combined the read counts corresponding to ASVs annotated as arbuscular mycorrhiza using FUNGuild and transformed them relative to the reads of all other eukaryotic taxa identified with the marker. Differences between regions were tested with one-way nested ANOVA or GLS models as described above. Pairwise differences were tested with the Tukey test. Benjamini-Hochberg multiple testing correction was applied to *P*-values at a false discovery rate of 0.05.

To show how often ASVs and taxa that are shared among or are unique to different groups, we used non-rarefied read counts with a relative abundance threshold of 0.001 to remove only the least abundant ASVs or taxa. We also checked the frequency of taxa without filtering for relative abundance and obtained very similar results. Taxa were classified as present within a specific group (genetic group, region, site), if at least one sample in that group had the given taxon.

## Results

### Soil properties contribute to the diversity of soil microbial communities in aspen-dominated stands

We compared soil and microbiome characteristics between two genetic groups: Canada and Mexico (NENA and MX in (Goessen *et al*. 2022)), and across three ecological regions—boreal, cold temperate, and warm temperate—referred to as regions. Soils from the two genetic groups were distinguished based on soil physicochemical properties (Fig. 1D-E, Supplementary Tables 4-5). Nested ANOVA analyses showed that the strongest differences between genetic groups included higher Fe in Canadian soils and higher K and Mn in Mexican soils (Supplementary Fig. 2). Other properties varied primarily between sites and/or regions. For example, Mexican (warm temperate) soils had on average higher pH, but the levels between sites varied considerably. Boreal sites were characterized by the highest levels of Mg (along with Fe), whereas cold temperate sites showed the lowest levels of Mn, Mg, K, and Ca. Macronutrients (N, C, and P) did not vary between genetic groups, but mostly between sites and regions. Boreal sites scored the highest levels of N and C but had the lowest C:N ratio. Boreal and warm temperate sites also showed high levels of cation exchange capacity (CEC) (Supplementary Fig. 2).

Bacterial, archaeal, and fungal communities were distinct between the genetic groups, regions, and between sites (PERMANOVA, *P* < 0.001, Fig. 2A-B). We investigated the impact of soil properties and genetic relatedness of trees on microbial community structure. Community structure was correlated with many soil properties (envfit analysis, *P* < 0.05, Fig. 2A-B, Supplementary Table 6), suggesting that they were driving part of the variation of bacterial, archaeal, and fungal communities. In bacteria and Archaea, the most correlated properties included clay and sand texture, Fe, Ca, and Mg (*R*^2^ = 0.5-0.7, Supplementary Table 6), whereas in fungal communities it was mainly K, Mn, and Ca (*R*^2^ = 0.44-0.48, Supplementary Table 6).

**Fig 2.**
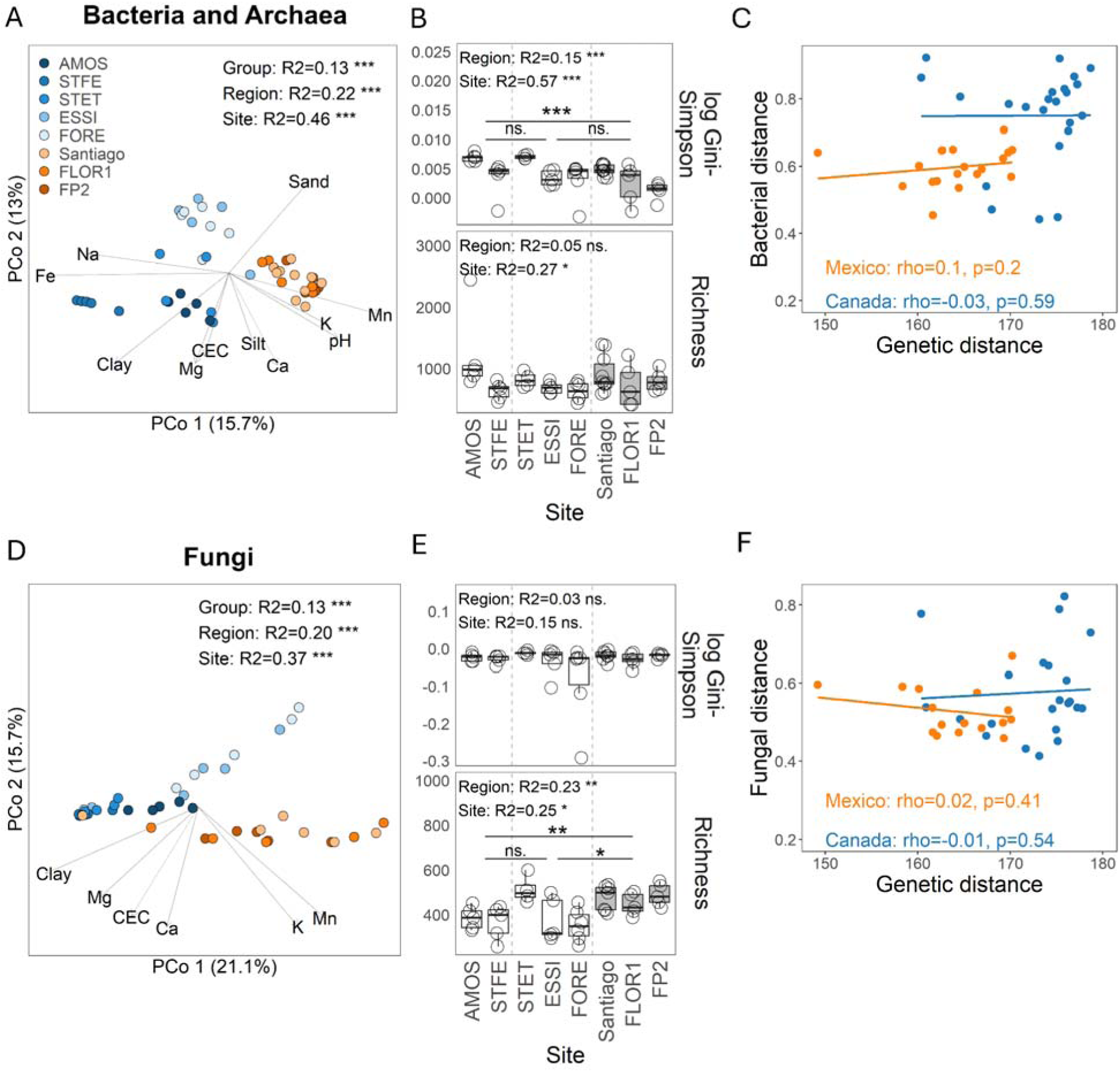
Factors driving soil bacterial, archaeal, and fungal community structure. Ordination of A) bacterial/archaeal, and D) fungal communities. Arrows show significant soil properties fitted onto the ordination of microbial structure with the envfit analysis. Only soil properties with significant vectors at *P* < 0.01 are shown. *R^2^*values refer to PERMANOVA analysis for each grouping level: genetic group, region or site. B, E) Alpha diversity metrics in bacterial and archaeal (B) and fungal (E) communities. Differences between regions and sites were tested with nested ANOVA, and pairwise differences with the Tukey test. C, F) Mantel test comparing dissimilarities of bacterial and archaeal (C) and fungal (F) community compositions with distances of individual tree genotypes. Principal component analysis on aspen variants is shown in Supplementary Figure S3.

Because at the global scale, soil differences can be correlated with other factors specific to genetic groups, we also investigated the impact of soil at the smaller, genetic group scale. In Canada, bacterial/archaeal community structure was mostly correlated with Ca, Mg and clay (*R*^2^ = 0.75-0.88, Supplementary Table 7), and fungal communities with Ca, Mn and CEC (*R*^2^ = 0.41-0.46, Supplementary Table 7). In Mexico, bacterial/archaeal communities were best correlated with silt, pH, and CEC (*R*^2^ = 0.6-0.71, Supplementary Table 8), whereas for fungal communities Fe was the dominating factor (*R*^2^ = 0.36, Supplementary Table 8).

To measure the impact of aspen genetic variation on the microbial community structure, we compared genetic distances between individual trees sequenced in a previous study (Supplementary Fig. 3, (Goessen *et al*. 2022)) with the dissimilarities between soil microbiome community compositions collected under those trees. As the Mexican and Canadian aspens belong to two distinct genetic groups, we measured the distances within each genetic group, to avoid confusion of genetic differences with other factors. We found that the aspen genetic distance was not correlated with the microbial community dissimilarities (Mantel test, *P* < 0.05, Fig. 2C,G).

Alpha diversity indices varied regionally for bacterial and archaeal communities (Gini-Simpson index, *F* = 9.7, *P* = 4.13E-04, Fig. 2B), and for fungal communities (richness, *F* = 7.7, *P* = 0.002, Fig. 2F). Boreal sites had a higher bacterial/archaeal diversity than warm temperate sites (Gini-Simpson index, Tukey test, *P* = 3.14E-04), whereas warm temperate sites showed higher fungal richness than both boreal and cold temperate sites (Tukey test, *P* < 0.02). However, alpha diversity metrics varied even more between sites (Fig. 2B,F). We tested if soil nutrients were driving variation in alpha diversity. As many soil and climate factors influence diversity of bacterial and fungal groups (Labouyrie *et al*. 2023), we modeled Gini-Simpson index using beta regression, with the following uncorrelated variables: pH, C:N, P, K, and Na, and a group as a fixed effect to control for multiple factors varying between the Canadian and Mexican samples. In bacteria/Archaea, the best model explained a large portion of diversity (pseudo *R*^2^ = 0.58, Supplementary Table 9). The main predictors included interactions between the group, pH and C:N, and the overall effect of Na. The model revealed negative correlations between diversity and pH, or diversity and C:N in Mexico but not in Canada (Supplementary Table 9). In fungi, neither the selected soil variables nor the group had a substantial effect on diversity (pseudo *R*^2^ = 0.12, Supplementary Table 9). Alpha diversity of bacteria/Archaea and fungi were positively correlated with each other (Spearman’s Rank correlation, rho = 0.38, *P* = 0.01).

We observed up to 3 times log-fold changes in the taxonomic composition of microbial taxa among boreal and temperate regions (genus level, Fig. 3, family level, Supplementary Fig. 4).

**Fig 3.**
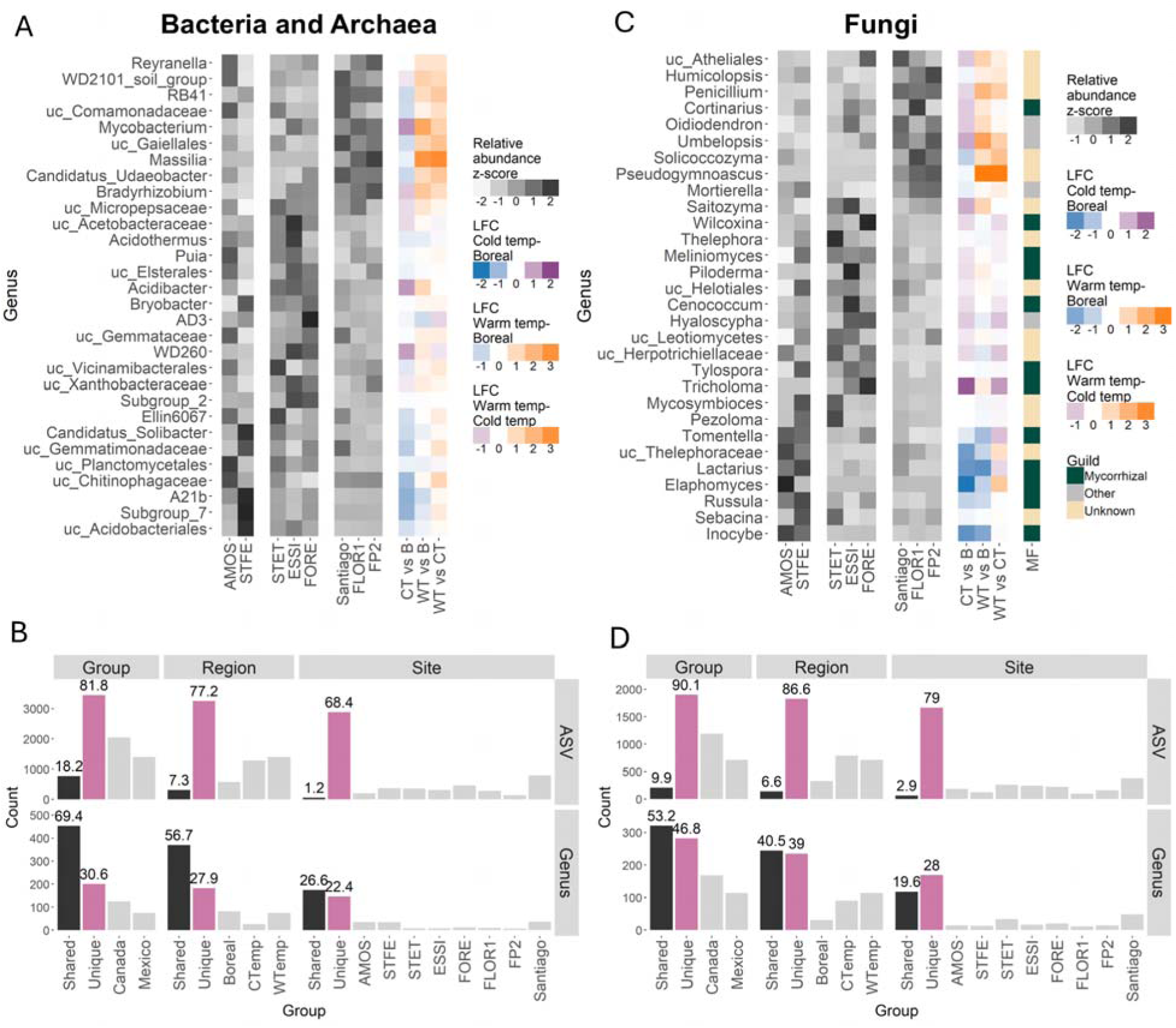
Taxonomic composition of the most highly abundant soil taxa. A, C) Grey heatmaps show standardized relative abundances and colored heatmaps show differential abundance (natural log fold change, LFC) between the pairs of regions in bacterial and archaeal (A), and fungal (C) taxa clustered at the genus level. Taxa most abundant in boreal region (B) are in blue, in cold temperate region (CT) in purple, and in warm temperate region (WT) in orange. The second color heatmap in (C) named “MF” indicates taxa classified as mycorrhizal fungi (green), other guild (grey), or unknown (beige) using FUNGuild. Prefix “uc” in taxonomic names indicates all unidentified or uncultured genera clustered at the higher level. Genera are ordered such that the topmost genera have the highest mean relative abundance in the warm temperate region, the middle ones - in the cold temperate region, and the bottom ones - in the boreal region. B, D) Counts of ASVs and genera shared across (black), or unique to genetic group, region, or site (pink), for bacterial and Archaea (B), and fungi (D). Grey bars indicate the number of taxa specific to the given class. Note that categories are not mutually exclusive. The numbers above bars indicate percentages of all taxa. Differential abundance between bacterial, archaeal, and fungal families is shown in Supplementary Figure S4.

However, none of the highly abundant genera showed significant differences in relative abundance between the pairs of regions, likely due to high variability across sites (Fig. 3A, C). At the genus level, a large proportion of bacterial and archaeal microbiome was shared between Canada and Mexico (69.4%, Fig. 3B) or between the three regions (56.7%, Fig. 3B), whereas much fewer genera were unique to a genetic group or a region (30.6% and 27.9%, respectively, Fig. 3B). In fungi, the number of genera unique to genetic group or region (46.8% and 39% respectively, Fig. 3D), was similar to the number of genera shared by the groups (53.2% and 40.5%, respectively, Fig. 3D), indicating higher regional specificity of fungal genera compared to bacteria. Conversely, at the ASV level, most variants were unique to the genetic group, region, or site (Fig. 3B, D). Bacterial genera present across all sampled sites primarily belonged to the Phyla *Proteobacteria* (n = 52), *Acidobacteriota* (n = 21), *Chloroflexi* (n = 18), *Actinobacteriota* (n = 17), *Planctomycetota* (n = 16), and *Bacteroidota* (n = 14), among others. Shared fungal genera were mostly from *Ascomycota* (n = 66) and *Basidiomycota* (n = 44), with smaller contributions from *Mucoromycota* (n = 3), *Rozellomycota* (n = 2), *Chytridomycota* (n = 1), and *Mortierellomycota* (n = 1).

Boreal and warm temperate regions were taxonomically most differentiated (Fig. 3, Supplementary Fig. 4). Bacterial taxa most enriched in the boreal region included *Ktedonobacteracea*, A21b, and *Subgroup 7*, whereas the warm temperate region of Mexico had highly abundant families such as *Oxalobacteraceae* (*Massilia* spp.), *Mycobacteriaceae* (*Mycobacterium* spp.), or *Burkholderiaceae*. Cold temperate sites had only a few taxa that were more abundant than in both boreal and warm temperate sites (WD260), with most taxa having a relative abundance intermediate between the two regions (*Ktenobacteracea*, AD3, *Mycobacteracea*). Fungal taxa enriched in boreal sites included families *Elaphomycetacea* (*Elaphomyces* sp.), *Inocybaceae* (*Inocybe* spp.), and *Russulaceae* (*Lactarius* spp., *Russula* spp.), in cold temperate region *Tricholomatacea* (*Tricholoma* sp.), and in the warm temperate region *Pseudeurotiaceae* (*Pseudogymnoascus* spp.), *Aspergillaceae* (*Penicillium* spp.), and *Umbelopsidaceae* (*Umbelopsis* spp.). Notably, fungal genera with higher abundances in boreal and cold temperate sites were often ectomycorrhizal (Fig. 3B).

### Communities of ectomycorrhizal (EMF) and arbuscular mycorrhizal fungi (AMF) vary between ecozones

We compared the relative abundance of eight fungal guilds between boreal, cold, and warm temperate regions. Consistent with the prevalence of mycorrhizal fungi among the most enriched genera in Canada (Fig. 3C), we observed significant regional differentiation in ectomycorrhizal fungi (EMF, one-way nested ANOVA, *F* = 8.7, *P* = 0.002, Fig. 4A), the most abundant among eight analyzed fungal guilds (Supplementary Fig. 5). The boreal region had a higher relative abundance of EMF compared to both cold and warm temperate regions (Tukey test, *P* < 0.05), whereas cold and warm temperate regions had similar levels of EMF (Tukey test, *P* = 0.42). In contrast, alpha diversity remained constant across the three regions (*F* = 0.12, and *F* = 2.33, respectively for the Gini-Simpson index and species richness, *P* > 0.1, Fig. 4A).

**Fig 4.**
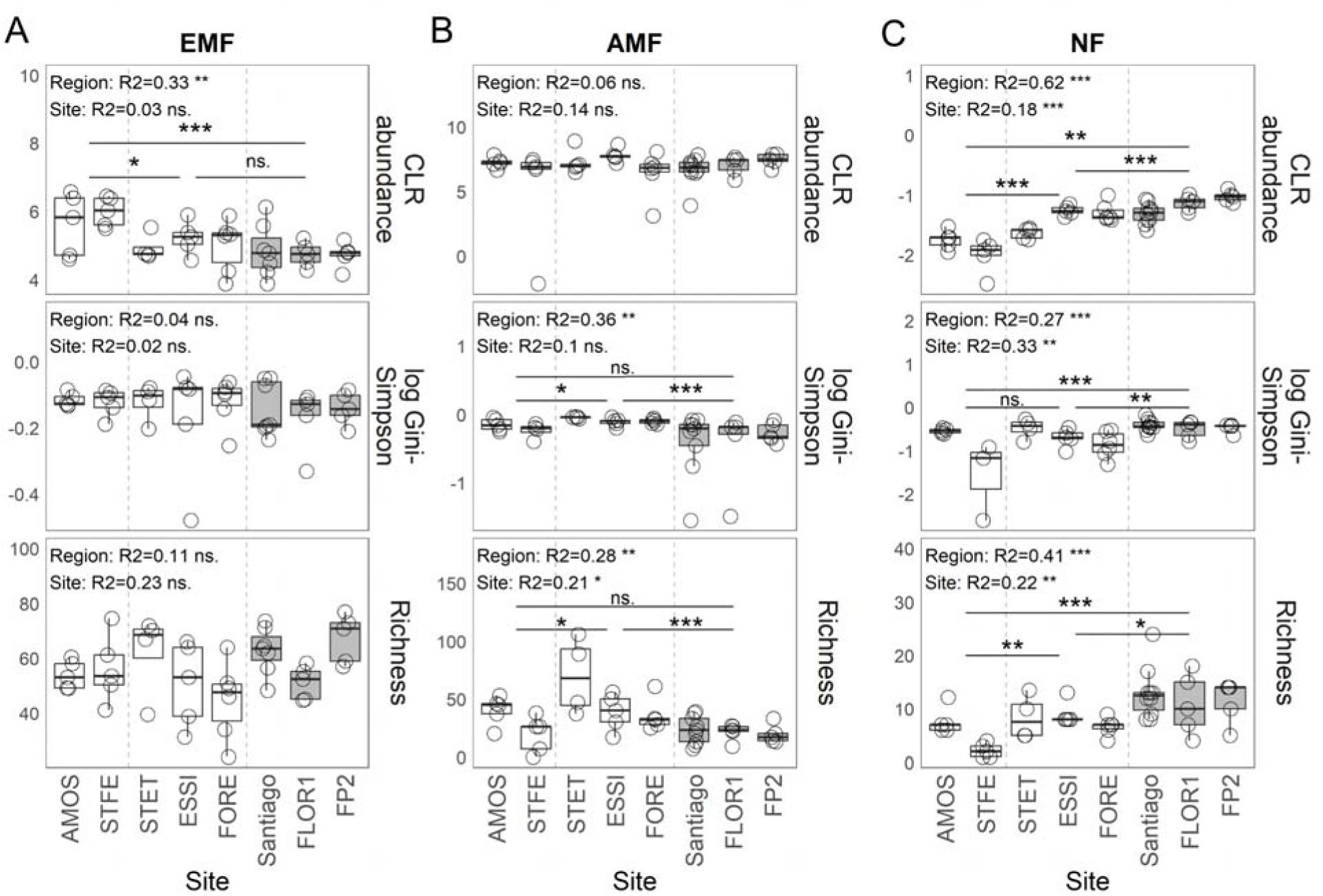
Abundance and alpha diversity of EMF (A), AMF (B), and selected nitrogen-fixing bacteria, NF (C) between the three regions of aspen natural stands. Sites are divided with dashed lines into boreal, cold temperate, and warm temperate regions. CLR stands for center log-ratio transformed abundance. Grey color indicates genetic groups or regions from Mexico. Differences between regions were tested using nested ANOVA. Pairwise differences between regions were performed with the Tukey test. ns. stands for not significant after multiple test correction with the Benjamini-Hochberg procedure at 0.05; * *P* < 0.05, ** *P* < 0.01,*** *P* < 0.001. Three Gini-Simpson outlier values were removed from the plot for better visibility.

Because the ITS2 region is not reliable in detecting highly variable rDNA regions of arbuscular mycorrhizal fungi (AMF), we sequenced all samples for the 18S rRNA gene using primers designed specifically to amplify AMF sequences. Since not only AMF but also other taxonomic groups were identified in our data set with 18S rRNA gene primers, we used the proportion of AMF reads relative to all reads captured by 18S rRNA gene as a proxy for their relative abundance. We found no difference in AMF relative abundance between the three regions (*F* = 1.24, *P* = 0.34, Fig. 4B). However, the Gini-Simpson index and richness differed significantly between the regions (*F* = 10.67, and *F* = 9.68, respectively, *P* = 0.001, Fig. 4B), with the highest levels found in the cold temperate region (Tukey test, *P* < 0.04, Fig. 4D).

To examine the relative abundance and alpha diversity of nitrogen-fixing bacteria in aspen stands, we searched our dataset for both symbiotic and free-living bacteria commonly known to carry out nitrogen fixation. This approach is not sufficient to identify all nitrogen-fixing ASVs but rather compares a selected group of taxa capable of this process mostly composed of *Frankia*, rhizobia and Cyanobacteria. (Sepp *et al*. 2023). We found significant differences between the regions in both relative abundance (*F* = 60.11, *P* = 3.06E-13, Fig. 4C) and alpha diversity (*F* = 11.95, *F* = 20.21, for Gini-Simpson index and richness, respectively, *P* < 5.0E-4, Fig. 4C). While samples from Mexico showed reduced relative abundance in EMF and lower diversity of AMF, both of which facilitate nitrogen acquisition, they also showed higher relative abundance of selected nitrogen-fixing bacteria than in the cold temperate region (Tukey test, *P* = 0.002) and boreal region (Tukey test, *P* = 6.48E-13). Similarly, alpha diversity indices were higher in Mexico than in cold temperate (Tukey test, *P* < 0.02, Fig. 4C) and boreal regions (Tukey test, *P* < 5.0E-4, Fig. 4C).

### Greenhouse common garden confirms the influence of the aspen genetic makeup on bacterial, fungal, and EMF communities

To disentangle the effects of substrate and host genotype on the community structure and EMF/AMF abundance, we characterized the microbiome in the potting mix, rhizosphere, and roots of aspen seedlings from 4 sites and 2 genetic groups (2 sites from Mexico, and 2 sites from Canada) grown in the same substrate and under the same ambient conditions (Fig. 1C). We found that community structures differed between the sample types of bacterial/archaeal (PERMANOVA, *R*^2^ = 0.19, *P* < 0.01, Fig. 5A) and fungal communities (*R*^2^ = 0.1, *P* < 0.01, Fig. 5B). Similar to results in the natural stands, we found significant differences between the two genetic groups(bacteria: *R*^2^ = 0.07, *P* < 0.01; fungi: *R*^2^ = 0.12, *P* < 0.01, Fig. 5A-B), and between sites (bacteria: *R*^2^ = 0.17, *P* < 0.01; fungi: *R*^2^ = 0.23, *P* < 0.01, Fig. 5A-B) When analysing separately communities from the potting mix, rhizosphere or the roots, we observed larger differentiation of all communities between sites than genetic groups in all sample types (Fig. 5, Supplementary Figs. 6-7), but the overall effects were non-significant in most cases given a small sample size. These results confirmed that community differences were influenced by the genetic variation of the host either between or within the groups. No particular genus dominated in either of the genetic groups (Fig. 5C-D, Supplementary Figs. S6-S7). Alpha diversity indices in the potting mix, rhizosphere, and root samples were similar between the genetic groups and sites (Fig. 5E-F, Supplementary Figs. S6-S7).

**Fig 5.**
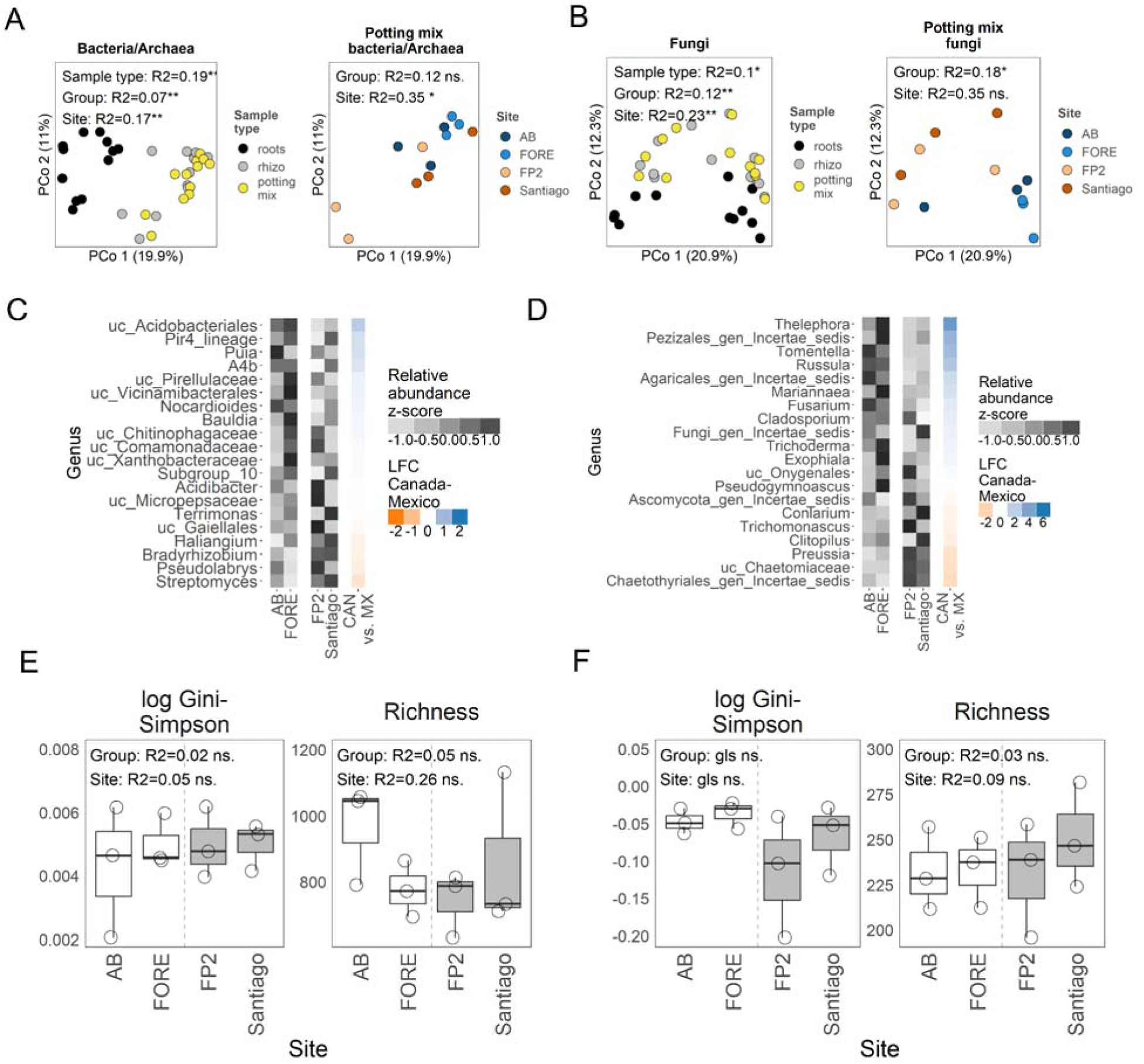
Communities from the potting mix of aspen seedlings from Canada and Mexico growing in greenhouse conditions. A-B) Ordination and PERMANOVA analysis of bacterial/archaeal (A) and fungal (B) communities visualized for samples from the three sample types (potting mix, rhizosphere, roots) and for potting mix samples only. C-D) Grey heatmaps with standardized relative abundances and colored heatmaps with differential abundance (natural log fold change, LFC) between the genetic groups in bacterial/archaeal (C), and fungal (D) taxa clustered at the genus level. Taxa differentially abundant in Mexico are in orange, and differentially abundant in Canada are in blue. E-F) Comparison of alpha diversity for bacterial/archaeal (E), and fungal (F) communities. Grey boxes indicate samples from Mexico. Differences between genetic groups were tested using nested ANOVA or generalized least squares (gls). ns. stands for not significant after multiple test correction with Benjamini-Hochberg procedure at 0.05; * *P* < 0.05, ** *P* < 0.01, *** *P* < 0.001. Results for roots and rhizosphere communities are shown in Supplementary Figures S6-S7.

We examined whether aspen from Canada forms more associations with mycorrhizal fungi than those from Mexico and whether Mexican aspen promotes more nitrogen-fixing bacteria than their Canadian counterparts when grown under the same conditions. We did not find AMF in any of the samples, likely because they were absent or not able to develop in the potting mix.

Regarding EMF, we found in total six genera in the potting mix, rhizosphere, and roots (Fig. 6B-D), which together had higher relative abundance in aspen plants from Canada in all three sample types (generalized least squares, *P* = 3.86E-07, and *P* = 1.26E-12; nested ANOVA, *P* = 0.07, for the potting mix, rhizosphere and roots respectively, Fig. 6A). This is in contrast to other fungal guilds which had similar abundances between the two genetic groups (Supplementary Fig. S8). Notably, the six EMF genera were present in both natural stand sites, and the roots of aspen seedlings (except for the genus *Laccaria*), indicating that they can form symbiotic associations with aspen. No significant differences in alpha diversity were found between sites from the two genetic groups (Fig. 6A). We also found no differences in the relative abundance or alpha diversity of nitrogen-fixing bacteria between the two genetic groups (Supplementary Fig. 9), which could be a consequence of frequent fertilization of the potting mix. In general, we confirmed that EMF fungi were more commonly found in association with aspen from Canadian sites, than from Mexico.

**Fig 6.**
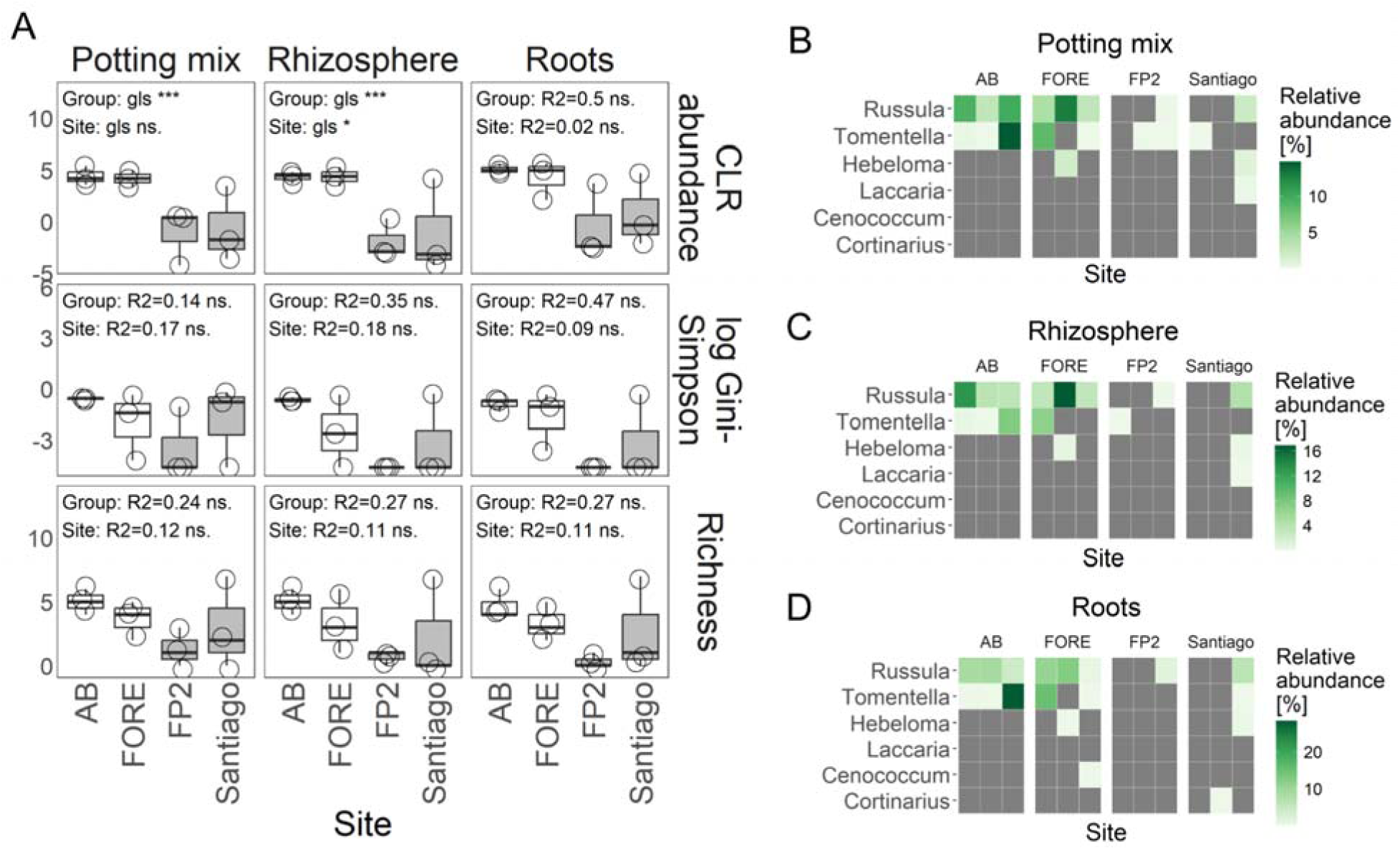
Relative abundance and alpha diversity of EMF communities of aspen seedlings from Canada and Mexico growing in greenhouse conditions. A) Comparison of center log-ratio transformed (CLR) abundance, log of Gini-Simpson index, and species richness in EMF communities between two genetic groups. Grey boxes indicate samples originating from Mexico. Differences between genetic groups were tested using nested ANOVA or generalized least squares (gls). B-D) Heatmaps with a relative abundance of EMF genera measured in potting mix(B), rhizosphere (C), and roots (D).

## Discussion

We investigated the soil communities associated with two genetically and geo-climatically distinct genetic groups of *P. tremuloides*: one sourced from Northwestern Mexico and the other from Eastern Canada. Our study examined how soil properties and host tree genotype influence the composition, structure and diversity of these communities and tested the impact of tree origin using a common garden setup. We found that both host genotype and soil properties contribute to the divergence of the aspen soil microbiome. Additionally, mirroring the geo-climatic variation across sampled sites, we observed differences in community structure and composition among three forest regions—boreal, cold temperate, and warm temperate. These regions were primarily differentiated by the relative abundance and diversity of ectomycorrhizal fungi (EMF), arbuscular mycorrhizal fungi (AMF), and nitrogen-fixing bacteria.

### Factors driving differences in diversity and community structure of aspen

The two studied aspen genetic groups diverged around 800,000 ya (Goessen 2023) and occupy somewhat distinct habitats at the southern and northern extremes of the species’ range: Northwestern Mexico and Quebec, Canada, respectively. Due to the high elevation of the Mexican stands, this temperate genetic group experiences much cooler weather than the adjacent dry tropical forests and obtains nearly as much annual rainfall as temperate parts of Canada. Nevertheless, aspen stands from Mexico differ markedly from Canadian stands in lower temperature seasonality and a different precipitation regime which is constrained to the rainy season. Other factors also vary between the habitats of two genetic groups including abiotic (soil, altitude, photoperiod) and biotic (host genetic differentiation, forest taxonomic composition, and surrounding understory vegetation) – all of which can contribute to divergence in their associated microbiomes (Adair and Douglas 2017; Baldrian 2017; Guo *et al*. 2023; Chamard *et al*. 2024). We found that microbial communities indeed varied between the two genetic groups, however most variation was distributed among samples and among sites. This is consistent with findings of strong spatial variability and a smaller impact of seasonal changes on *P. tremuloides* bacterial and fungal root communities in the U.S.A. (Nash *et al*. 2025). In our study, we focused on analyzing the impact of soil properties and host genotype on the overall microbial community.

Soil properties are one of the key factors shaping microbiota, influencing microbial growth rates, diversity and composition primarily through pH and nutrient availability, and their interaction with climate (Rousk *et al*. 2010; Ding, Travers and Eldridge 2022; Sridhar *et al*. 2022; Osburn *et al*. 2024). For example, interaction between pH and temperature was shown to predict continental and global extent of forest soil bacterial richness much better than pH alone (Luan *et al*. 2023). We observed that pH varied only slightly between Mexico and Canada and levels of macronutrients such as N, C, P and remained similar, but they all varied significantly among the sites. However, the overall combination of soil parameters clearly differentiated the two genetic groups. At the continental scale, Fe, K, Ca, Mn correlated best with microbial structure, but they also strongly differed between groups, making it difficult to disentangle the effect of those nutrients from other group-specific factors. Nevertheless, several nutrients were strongly correlated with community structures (*R*^2^ = 0.5-0.9) also on the local (genetic group) levels, suggesting at least partial contribution of soil to aspen microbiome differences between sites and regions. We also found that a part of bacterial diversity was explained by the interactions between the group, pH and C:N. While we could not measure the effects of climate variables such as temperature, humidity or photoperiod on these variables, the fact that the effects of soil variables on diversity differed among the groups suggests that local geo-climatic conditions and/or host genetic group influence the way microbes respond to soil conditions. The year of sampling is another factor that could potentially influence diversity analyses (Dove *et al*. 2021). All sites in Mexico were sampled in 2019 and all sites in Canada except for one (STET sampled in 2019) were sampled in 2018, which makes it impossible to disentangle the effects of sampling from other group-related effects. Within the Canada group, the Gini-Simpson index values for the only site sampled in a different year match closely the distribution of values estimated across other sites. While our results suggest that inter-year differences in diversity do not exceed inter-site differences, we cannot exclude that some of our results are driven by temporal rather than geographic factors.

Host plant genotypes can influence soil microbiomes through root exudates that attract and inhibit specific microbes (Morgan, Bending and White 2005). Metabolic profiles of root exudates show natural variation between (Rathore *et al*. 2023) and within species (Mönchgesang *et al*. 2016) that eventually impact the composition of the rhizosphere microbiome (Seitz *et al*. 2022). Aspen is genetically structured into four major genetic groups, with those from Mexico and Eastern Canada being the most divergent (*F*_ST_ = 0.174; (Goessen *et al*. 2022)). Within genetic groups, genetic differentiation between sites is low, accounting for just 0.3–4.4% of the total variation (Goessen *et al*. 2022). Consistent with this, we found no relationship between aspen genotypes and their microbial communities within genetic groups of natural stands. The strongest trend was observed in bacteria and Archaea from Mexico. Aspen trees from Mexico are more differentiated, likely due to clonal reproduction in isolated mountainous areas, which could explain a stronger trend in this genetic group. This suggests that at the observed level of genetic differentiation, host genotype effects are overshadowed by other factors. Our results agree with previous common gardens conducted in *Populus* species, showing that soil physico-chemical properties, soil origin and host genotype all influence microbial communities, but the effects of soil are stronger, whereas the host genotype effects are not evident in every soil condition (Bonito *et al*. 2019; Veach *et al*. 2019; Rheault *et al*. 2020).

In contrast, greenhouse common garden experiments demonstrated the influence of genotype at both local (between study sites) and global (between genetic groups) scales. Although host genotype effects (group and site effects) are evident in spite of small sample size, two considerations should be taken into account. First, the seedlings were grown in the soil-less potting mix with an unknown source of microbes. In this study design, we cannot measure the effects the host genotypes have on soil microbiomes found at the tree’s site of origin and the effects on the soil microbiomes of foreign origin. Second, we cannot exclude the possibility that some portion of the microbiome has been transmitted with the roots of Mexican seedlings, which we were unable to propagate using cuttings. Although this might have inflated the observed differences between groups or sites, we note that we observed consistently stronger differences in microbial composition between sites than groups, and that the site differences were visible in seedlings of both Mexican and Canadian origin, which suggests that they were influenced by the host genotype. Nevertheless, the natural recommended follow-up to this finding would require reciprocal transplants of Canadian and Mexican cuttings in their native and foreign soil substrate to understand the extent of the host genotype influence.

Because plant roots influence microbial community assembly by attracting and promoting growth of certain microbes (Badri *et al*. 2009), the surrounding vegetation can significantly shape the soil microbiome of aspen stands. In Canada and in the U.S.A., *P. tremuloides* is not typically associated with high-diversity environments. As a pioneer species, it is more commonly found in early-successional habitats or areas recovering from disturbances, such as forest fires, logging, or agricultural abandonment (Pinno and Errington 2015, Swift and Ran 2012; Shinneman *et al*. 2013; Stefani *et al*. 2018), where competition for light, nutrients, and space is minimal. Consequently, its presence is often restricted to specific niches or transitional phases in forest succession. On the other hand, in Mexico it can be associated with diverse habitats (Simental-Rodríguez *et al*. 2014). To assess the relative impact of vegetation, as well as soil, and host genotype on soil microbiome communities in natural stands, a detailed plant inventory, and a broader sampling would be necessary.

### A gradient in taxonomic composition of aspen soil microbiome

Most (∼70%) of the bacterial microbiome could be found in both Canada and Mexico. Bacterial taxa commonly found across all sampling sites in our study belong to phyla that were previously consistently found as common components of poplar soil (*Proteobacteria*, *Acidobacteriota*, *Chloroflexi*, *Actinobacteriota*, *Planctomycetota*) or poplar roots (*Bacteroidota*, (Zanin *et al*. 2024)), and were also in general the most commonly found phyla in soil and rhizosphere (Ling, Wang and Kuzyakov 2022). Two genera we noted as highly abundant, *Bradyrhizobium* and *Mycobacterium,* were also reported previously (Zanin *et al*. 2024). One commonly found genus, enriched in Mexico, was *Massilia*, which was previously found to be a core component of the *Populus* phyllosphere (Dove *et al*. 2021; Liu *et al*. 2023). Although taxonomic enrichments between regions were not significant, likely due to large heterogeneity across sites, some trends in the distribution of taxa were evident. For example, *Massilia*, *Bradyrhizobium* and *Mycobacterium* dominated in warm temperate sites of Mexico, and boreal sites had higher abundance of Subgroup 7 (*Acidobacteriota*), and A21b (*Burkholderiales, Proteobacteria*).

*Bradyrhizobium* is a genus containing nitrogen-fixing bacteria, and *Mycobacterium* is a widely distributed decomposer in soils across forest ecosystems (Walsh *et al*. 2019). *Massilia* is a common rhizosphere component (Ofek, Hadar and Minz 2012), and interestingly, it was shown to exhibit high phosphatase activity in forest soil when growth of fungal hyphae was suppressed (Brooks *et al*. 2011). Subgroup 7 is not commonly reported, but was previously found, e.g., in agriculturally modified soils in Brazil (Navarrete *et al*. 2015; de Chaves *et al*. 2019), while A21b in soils from Antarctica (Krauze *et al*. 2021). It is not clear however if these species have any association with aspen and may simply be common components of the soil in that region.

Fungal taxa proved to have a more constrained distribution, with around half of the genera (∼53%) common across and half (∼47%) specific to only one of the two genetic groups. Also, 39% of fungal genera were found uniquely within one of three climate regions. This observation is consistent with higher global estimates of environmental specificity for fungi than bacteria (Loos *et al*. 2024). Regional endemism was notably found in poplar mycobiome and was explained by combination of soil, climate effects and geographical distance (Van Nuland *et al*. 2023). Several most abundant fungal genera in our dataset were reported previously in soil nearby poplar (*Cenococcum*, *Cortinarius*, *Inocybe*, *Lactarius*, *Mortierella*, *Tomentella, Tricholoma*) (Cripps and Miller 1993; Zanin *et al*. 2024). Similar to bacteria, fungal taxa were not significantly enriched in any of the three regions, but few of them showed some regional preferences. Boreal sites were preferentially occupied by *Elaphomyces*, *Inocybe* and *Lactarius* and cold temperate sites by *Tricholoma*. The only common mycorrhizal genus preferentially found in Mexico was *Cortinarius*, and three non-mycorrhizal highly enriched taxa included *Pseudogymnoascus*, *Penicillium* and *Umbelopsis*. All three genera are known to be associated with the rhizosphere (Summerbell 2005; Xia, Rufty and Shi 2021).

### Shifts in mycorrhizal communities along a latitudinal gradient

Ectomycorrhizal fungi (EMF) and arbuscular mycorrhizal fungi (AMF) differ in their capabilities to acquire nutrients. AMF, which are associated with nearly 80% of all land plants, rely on phosphorus (P) and nitrogen (N) released by saprotrophic microbes in the surrounding soil (Phillips, Brzostek and Midgley 2013). In contrast, EMF are primarily associated with trees and possess enzymes that allow them to directly access N and P from organic matter (Phillips, Brzostek and Midgley 2013). These decomposing capabilities give EMF a competitive advantage in boreal forests, where organic material tends to accumulate, and where EMF indeed dominate (Steidinger *et al*. 2019). In addition, the diversity of plants tends to be correlated with the diversity of mycorrhizal fungi (Fei *et al*. 2022), predicting a higher MF diversity at sites with rich surrounding vegetation, including other tree and understory species.

In aspen, an expected latitudinal shift in EMF is evident: EMF are significantly more abundant in boreal sites compared to temperate stands in Canada and in Mexico. Greenhouse common garden indicated that these EMF patterns are driven by specific aspen genotypes rather than soil conditions or vegetation differences. We note, however, that in this study we did not compare the relative abundances of EMF between Canadian and Mexican seedlings in their respective soils of origin, therefore we cannot determine to what extent these differences are limited by the presence or absence of specific set of fungal taxa in the potting mix or potential transfer of microbial inocula with the roots in the case of Mexican samples. The importance of AMF communities partially aligned with our expectations. While AMF were anticipated to be more prevalent in the temperate and southernmost sites of the aspen range (Steidinger *et al*. 2019), we observed a uniform relative abundance across all sampling locations, however we noted the highest diversity in temperate stands in Canada. As we were not able to measure AMF associations in the greenhouse study, it is less clear whether this uniformity in abundance reflects consistent AMF associations across aspen genetic groups, if it is a consequence of differences in understory vegetation among sites (Fei *et al*. 2022), in particular the sparse vegetation in high-altitude aspen habitats in Mexico, or other climate or soil factors (Van Aarle, Olsson and Söderström 2002).

Trees can also obtain nitrogen through symbiotic or free-living nitrogen-fixing bacteria and Archaea (Boyd and Peters 2013). Globally, rhizobial nitrogen fixers are more diverse in hot, stable, and seasonally humid regions (Sepp *et al*. 2023) and are more abundant in arid biomes (Steidinger *et al*. 2019). This aligns with our findings, where the highest relative abundance and richness of selected nitrogen-fixing bacteria were observed in Mexico. This pattern may also explain the lack of exceptionally high AMF diversity in this genetic group, suggesting that nitrogen-fixing bacteria either complement or dominate nitrogen acquisition in aspen.

### Aspen stand persistence and dynamics in the face of climate change

*P. tremuloides* likely originated in southern North America (Goessen 2023), and had to adapt to novel temperature regimes while it expanded northward, particularly to boreal regions. Notably, three genes that were identified to diverge across climate gradients are responsible for auxin biosynthesis and distribution in roots (Goessen 2023). Auxins are plant hormones that regulate plant growth and development (Gomes and Scortecci 2021), but are also known to influence arbuscular mycorrhiza colonization and promote their development (Gutjahr 2014). Auxin signalling has been also proposed to play a role in the formation of poplar-*Laccaria* ectomycorrhizal roots (Vayssières *et al*. 2015). These genes may therefore be key candidates for plant-fungal interactions that emerged during adaptation to novel climate conditions. It is however unclear how these plant-microbe interactions will change in response to rapid climate change, including an increasing drought frequency.

In agroforestry, understanding the stability and consequences of plant-soil interactions on host tree performance is one of the key questions, especially in the context of ongoing climate changes. There are at least two ways in which such interactions can be of potential importance: in increasing tree productivity, and in improving tree resilience to environmental stresses. Some studies highlight the role of specific genotype-microbiome interactions in promoting plant productivity, by demonstrating that plant genotypes modify their rhizosphere microbiome differently due to variation in nitrogen cycling processes, and in that way they influence the biomass of subsequent generations growing in conditioned soil (Przybylska *et al*. 2024). This aligns with the resurgent idea that organisms can influence their environments in a way to promote ecosystem resilience (Hunter 2018). On the other hand, experimental studies show that host responses to stressful events can be modulated by the soil microbiome. Soil transplant experiments demonstrated that plant soil microbiomes from different geographic origins can increase tree survival faced with various climate stresses (Allsup, George and Lankau 2023) or can affect plant phenology (Ware *et al*. 2021). This raises interesting questions as to whether the established genotype-microbiome interactions are more important for the tree survival and productivity in the long-term, or soil supplementation can be sufficient to withstand extreme climate events. Forestry practices such as assisted migration, which are designed to promote forest resilience to anticipated change in climate conditions, are expected to break such interactions, the consequences of which are not understood. Similarly, climate change is expected to affect microbiomes both directly (Knight *et al*. 2024), or through shifts in host distributions, changes in their phenological timing, and increasing mismatches between symbiotic partners (Classen *et al*. 2015). Therefore, understanding the contribution of plant-soil interactions on plant resilience to global changes is of increasing importance, and beneficial microbial communities should be conserved and harnessed to support the desired tree responses in forest management (Heuertz *et al*. 2023).

## Conclusions

Our study suggests that soil physicochemical properties and host genotype influence soil microbiomes of two genetic groups of *P. tremuloides* from Mexico and Canada. Our findings highlight the regional importance of nutrient-exchange symbioses in aspen stands: EMF dominate in the boreal forest, AMF are most diverse in temperate forests, and the selected groups of nitrogen-fixing genera abound in seasonally dry, warm regions, aligning closely with the expected global patterns of symbiosis distribution. Aspen seedling genotypes from Canadian sites grown in the greenhouse associated more commonly with EMF, implying that plant genetics has an effect on host-microbial interactions that mirror global ecosystem functional divergence. Further experiments are needed to identify microbial species that differentially associate with aspen genotypes from ecological regions and to uncover the heritability and molecular mechanisms driving these interactions. These insights will lay the groundwork for future management strategies in sustainable agriculture and forestry incorporating and utilizing microbiomes.

## Availability of Data and Materials

All raw sequence data generated in this study have been deposited in Sequence Read Archive (SRA) under project accession number PRJNA1230202. Code is available at https://github.com/aniafijarczyk/Fijarczyk_etal_2025-AspenMicrobiome.

## Acknowledgments

We want to thank Lyne Touchette (sampling natural stands, sampling and root propagation in the greenhouse), Mebarek Lamara and Annie DesRochers (sampling natural stands), Denis Lachance (sample processing, DNA extraction), Fanny Michaud and Olivier Jeffrey (soil sample processing), Serge Rousseau (soil analysis), Caroline Bourdon and Karelle Rheault (sampling of greenhouse experiment, sample processing and DNA extraction).

