## Supplementary Figures and Tables for "Aspen-associated soil microbiomes reveal different strategies for nitrogen acquisition across ecosystems in Mexico and Canada"

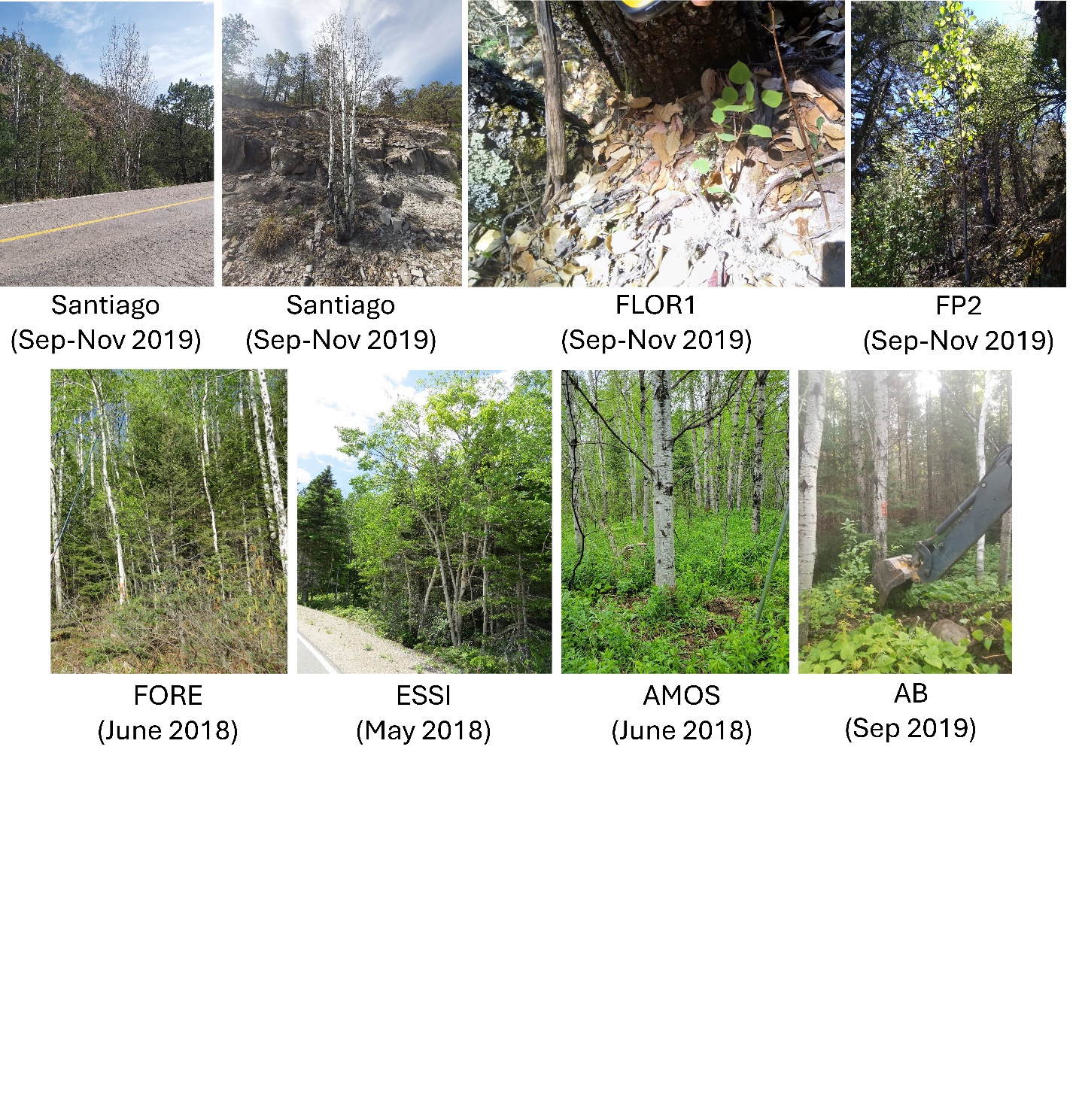

Fig S1. Diversity of sampled aspen habitats in Mexico (Santiago, FLOR1, FP2) and Canada (FORE, ESSI, AMOS, AB).

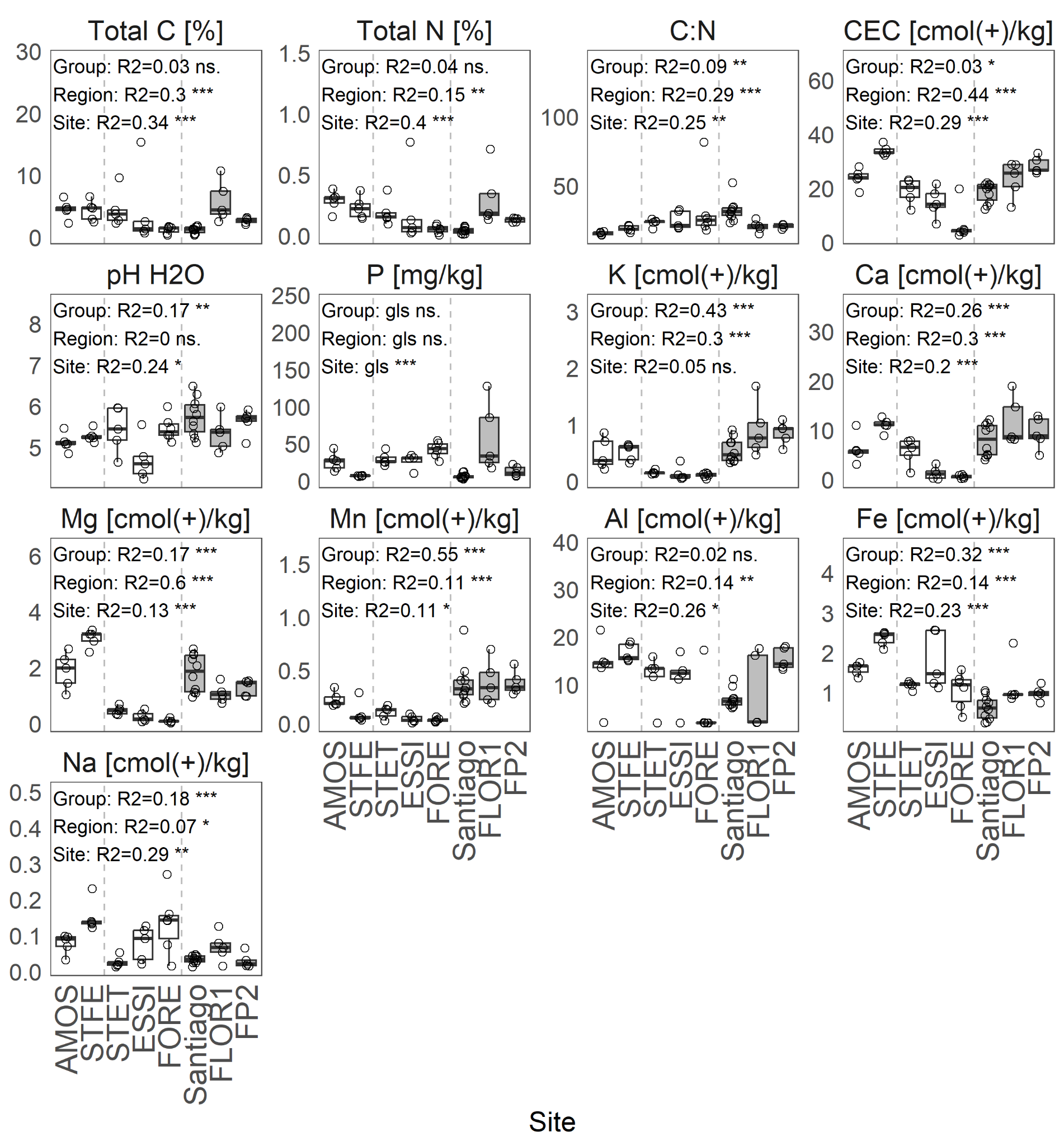

Fig S2. Comparison of soil physicochemical properties between genetic groups, regions and sites. Differences between genetic groups, regions and sites were tested using nested ANOVA or generalized least squares (gls) in case of non-homogeneity of variance. Grey boxes indicate samples originating from Mexico. Dashed lines separate three regions: boreal, cold temperate, and warm temperate. ns. stands for not significant after multiple test correction with Benjamini-Hochberg procedure at 0.05; * *P* < 0.05, ** *P* < 0.01, *** *P* < 0.001. A single phosphorus (P) outlier (P>400) was removed for better visibility.

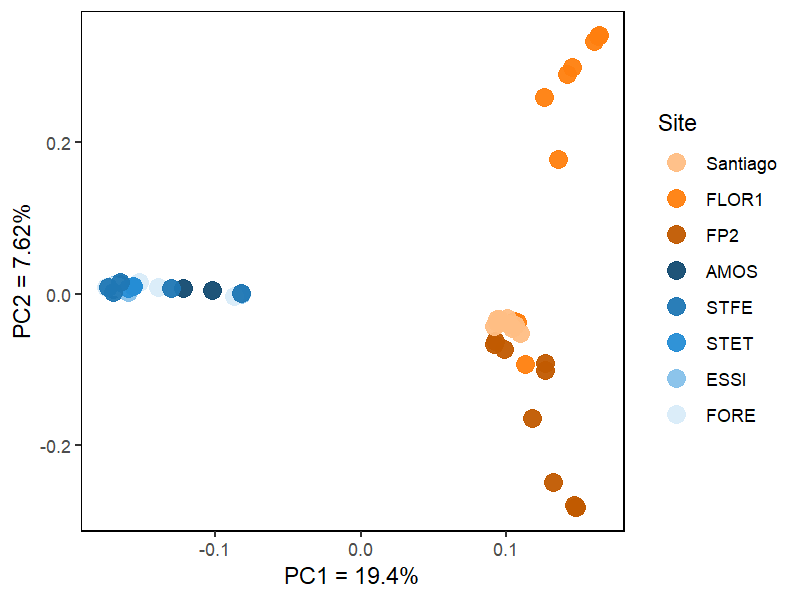

##### Fig S3. Principal component analysis using 53,714 genome-wide SNPs sequenced from aspen trees.

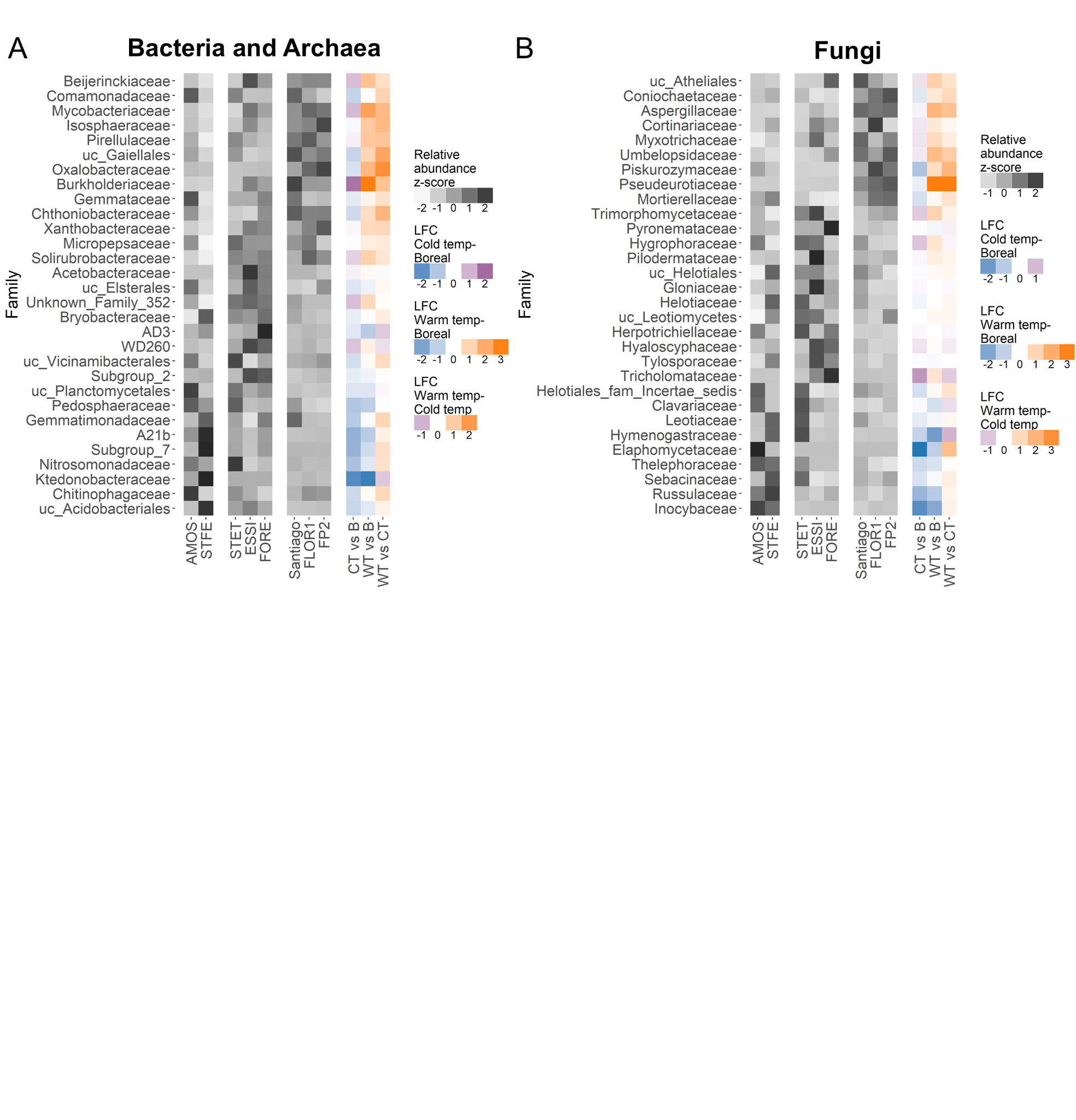

Fig S4. Differential abundance across three regions (boreal B, cold temperate CT and warm temperate WT) in top 30 most abundant soil families. A, B) Standardized relative abundances and differential abundances between the pairs of regions in bacterial and archaeal (A), and fungal (B) taxa clustered at the family level. Color heatmaps show estimates of natural log fold change (LFC) obtained with ANCOM-BC between pairs of regions. Taxa most abundant in boreal region (B) are in blue, in cold temperate region (CT) in purple, and in warm temperate region (WT) in orange. Prefix “uc” in taxonomic names indicates all unidentified or uncultured families clustered at the higher level. Genera are ordered such that topmost families have the highest mean relative abundance in warm temperate region, middle ones - in the cold temperate region, and bottom ones - in the boreal region.

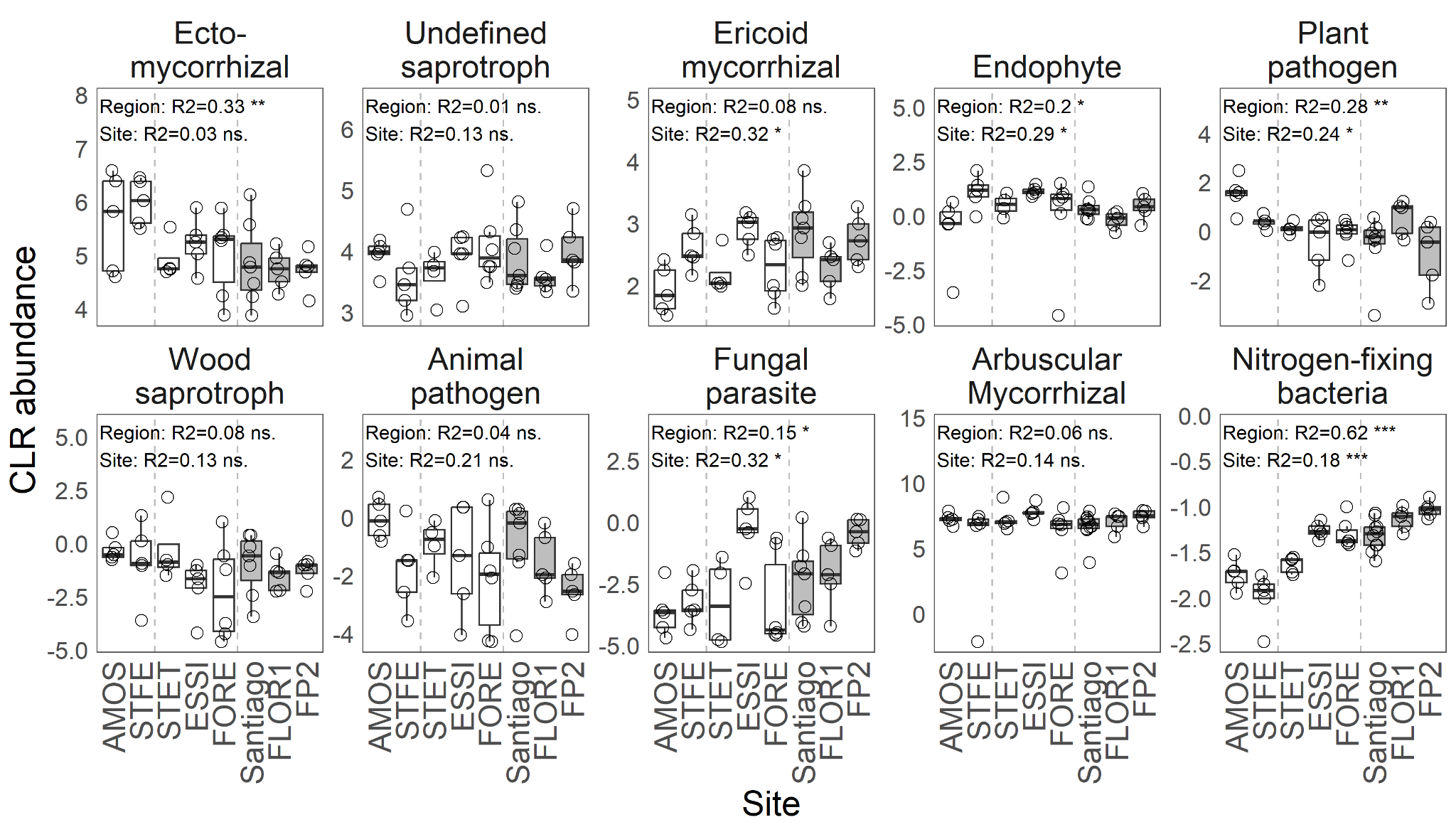

Fig S5. Comparison of log-ratio transformed (CLR) abundance of eight selected fungal guilds including EMF (ITS dataset), along arbuscular mycorrhiza, AMF (18S dataset), and selected nitrogen-fixing bacteria (16S dataset) between regions. Grey boxes indicate samples originating from Mexico. Dashed lines separate three regions: boreal, cold temperate, and warm temperate. Differences between genetic groups were tested using nested ANOVA. Ns. stands for not significant after multiple test correction with Benjamini-Hochberg procedure at 0.05; * *P* < 0.05, ** *P* < 0.01, *** *P* < 0.001.

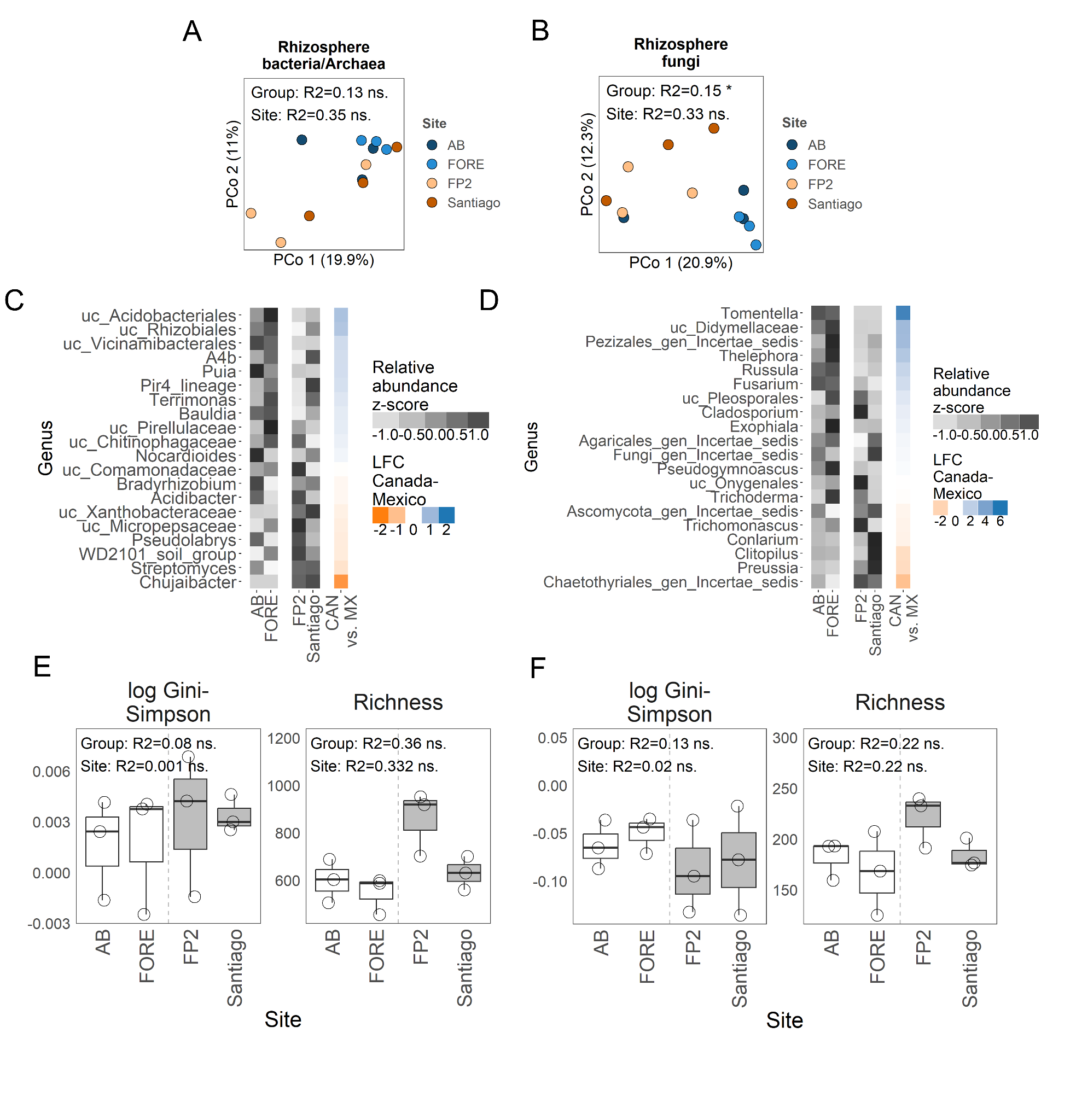

Fig S6. Rhizosphere communities of aspen seedlings from Canada and Mexico growing in greenhouse conditions. A-B) Ordination and PERMANOVA analysis of bacterial/archaeal (A) and fungal (B) communities. C-D) Grey heatmaps with standardized relative abundances and colored heatmaps with differential abundance (natural log fold change, LFC) between the genetic groups in bacterial/archaeal (C), and fungal (D) taxa clustered at the genus level. Taxa differentially abundant in Mexico are in orange, and differentially abundant in Canada are in blue. E-F) Comparison of alpha diversity for bacterial/archaeal (E), and fungal (F) communities. Grey boxes indicate samples from Mexico. Differences between genetic groups were tested using nested ANOVA. ns. stands for not significant after multiple test correction with Benjamini-Hochberg procedure at 0.05; * *P* < 0.05, ** *P* < 0.01, *** *P* < 0.001.

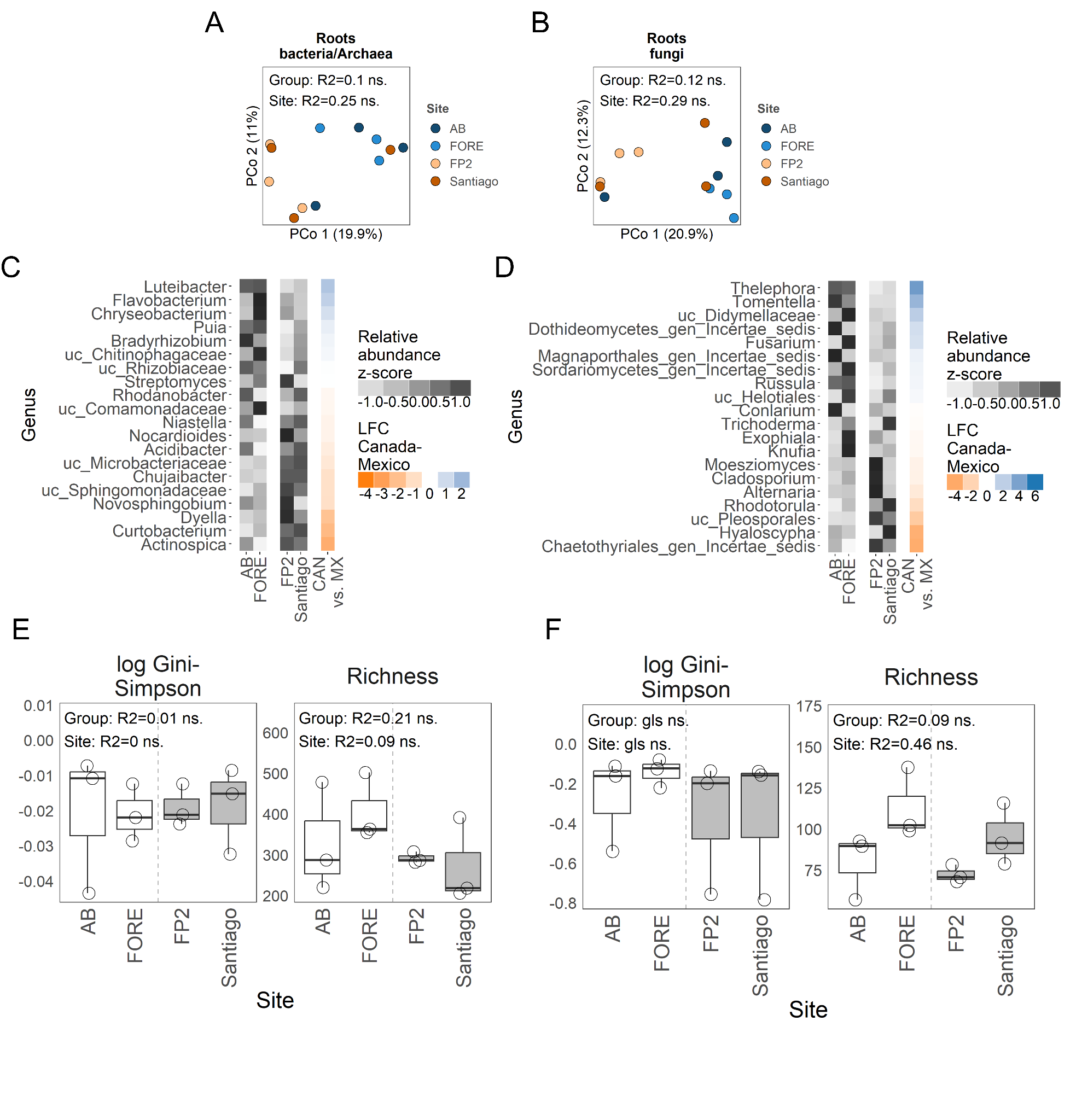

Fig S7. Roots communities of aspen seedlings from Canada and Mexico growing in greenhouse conditions. A-B) Ordination and PERMANOVA analysis of bacterial/archaeal (A) and fungal (B) communities. C-D) Grey heatmaps with standardized relative abundances and colored heatmaps with differential abundance (natural log fold change, LFC) between the genetic groups in bacterial/archaeal (C), and fungal (D) taxa clustered at the genus level. Taxa differentially abundant in Mexico are in orange, and differentially abundant in Canada are in blue. E-F) Comparison of alpha diversity for bacterial/archaeal (E), and fungal (F) communities. Grey boxes indicate samples from Mexico. Differences between genetic groups were tested using nested ANOVA or generalized least squares (gls). ns. stands for not significant after multiple test correction with Benjamini-Hochberg procedure at 0.05; * *P* < 0.05, ** *P* < 0.01, *** *P* < 0.001.

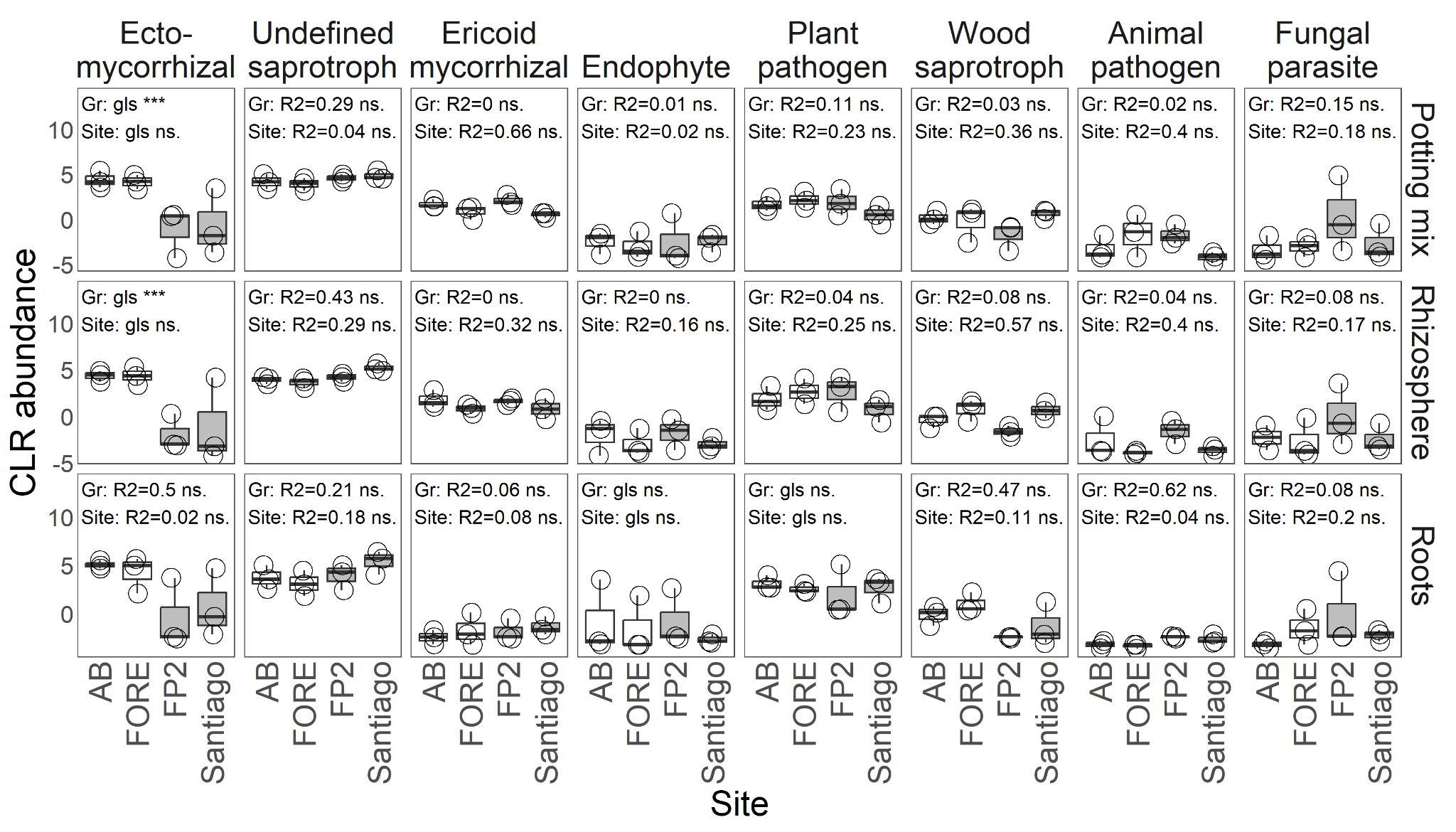

Fig S8. Comparison of log-ratio transformed (CLR) abundance of eight selected fungal guilds including EMF between genetic groups (Gr) in the potting mix, rhizosphere, and root samples from the greenhouse common garden. Grey boxes indicate samples originating from Mexico. Differences between genetic groups were tested using nested ANOVA or generalized least squares (gls). ns. stands for not significant after multiple test correction with Benjamini-Hochberg procedure at 0.05; * *P* < 0.05, ** *P* < 0.01, *** *P* < 0.001.

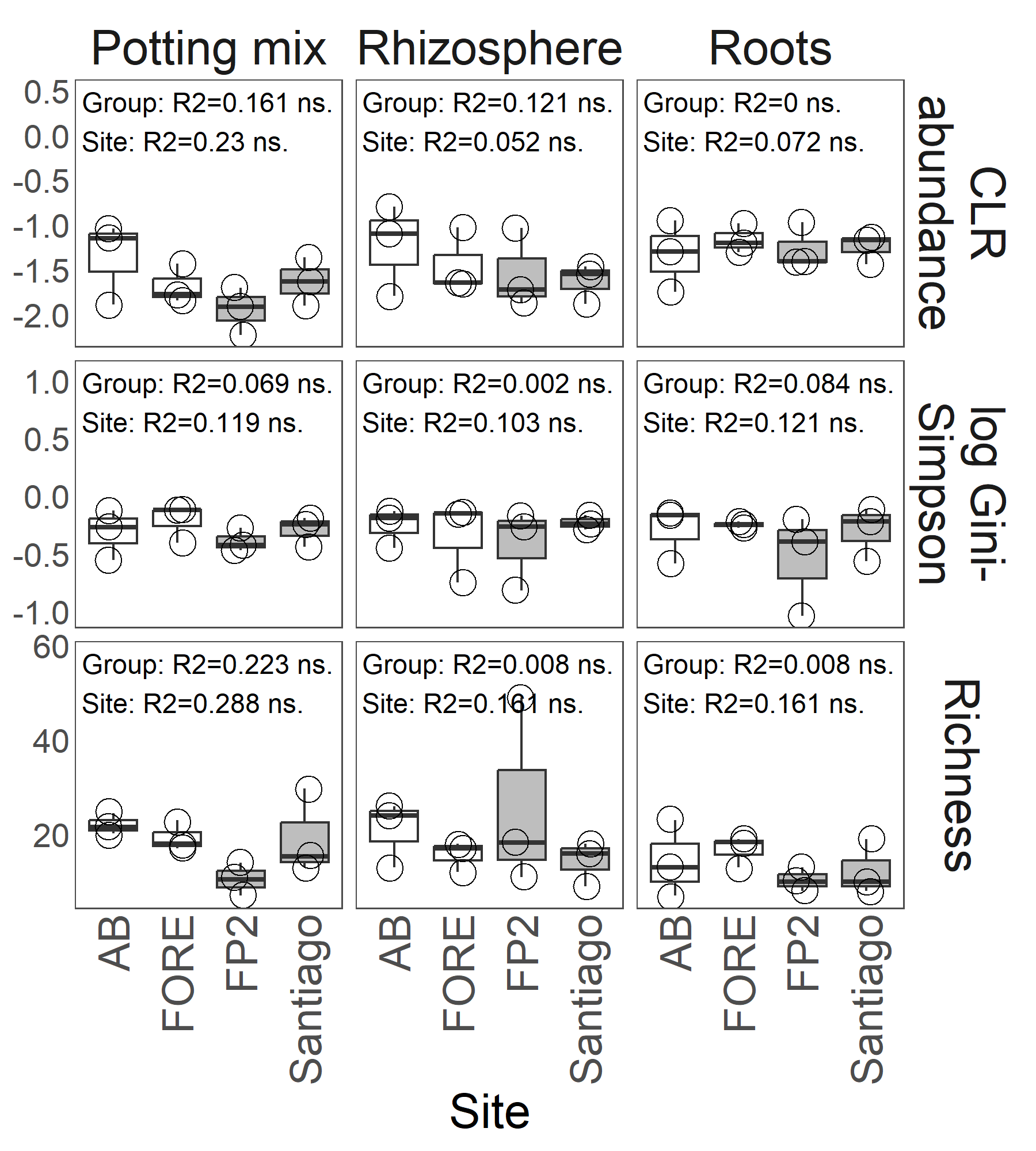

Fig S9. Comparison of log-ratio transformed (CLR) abundance, and alpha diversity of selected nitrogen-fixing bacteria between genetic groups in potting mix, rhizosphere, and root samples from the greenhouse experiment. Grey boxes indicate samples originating from Mexico. Differences between genetic groups were tested using nested ANOVA. ns. stands for not significant after multiple test correction with Benjamini-Hochberg procedure at 0.05.

Table S1. Description of sampled sites.

| **Site** | **Natural stand^1^** | **Green- house^1^** | **Year^2^** | **Group^3^** | **Group^4^** | **Adjacent town/village** | **Lat** | **Lon** |
| --- | --- | --- | --- | --- | --- | --- | --- | --- |
| AMOS | 5 | 0 | 2018 | Canada | NENA | Amos, Quebec | 48.616 | -78.073 |
| STFE | 5 | 0 | 2018 | Canada | NENA | St-Felix-De-Dalquier, Quebec | 48.710 | -78.116 |
| AB | 0 | 3 | 2019 | Canada | NENA |  | 48.300 | -79.220 |
| ESSI | 5 | 0 | 2018 | Canada | NENA | Essipit, Quebec | 48.341 | -69.400 |
| FORE | 6 | 3 | 2018 | Canada | NENA | Forestville, Quebec | 48.739 | -69.088 |
| STET | 5 | 0 | 2019 | Canada | NENA | St-Etienne-De-Lauzon, Quebec | 46.650 | -71.302 |
| Santiago | 10 | 3 | 2019 | Mexico | MX | Santiago | 25.028 | -105.702 |
| FP2 | 5 | 3 | 2019 | Mexico | MX | Aserradero-La-Flor | 23.524 | -104.685 |
| FLOR1 | 5 | 0 | 2019 | Mexico | MX | Aserradero-La-Flor | 23.531 | -104.691 |

^1^ The number of samples taken from natural stand site or greenhouse common garden

^2^ Year sample was taken (soil and/or root)

^3^ Genetic group

^4^ Genetic group code as in Goessen *et al*. 2022

Table S2. Description of basic soil and climate information of sampled sites.

| **Site** | **Texture** | **Sand [%]** | **Silt [%]** | **Clay [%]** | **Elev [m]^1^** | **MAT [C]^2^** | **Cont [C]^3^** | **MAP [mm]^4^** | **W&SP [mm]^5^** |
| --- | --- | --- | --- | --- | --- | --- | --- | --- | --- |
| AMOS | Clay | 12 | 34 | 54 | 317 | 1.6 | 33.9 | 872 | 313 |
| STFE | Heavy clay | 16 | 22 | 62 | 307 | 1.6 | 34 | 869 | 310 |
| AB | - |  |  |  | 344 | 2.1 | 34.1 | 860 | 324 |
| ESSI | Loamy sand | 83.99 | 8 | 8 | 42 | 3.5 | 29.3 | 1024 | 471 |
| FORE | Sandy loam | 74.09 | 17.94 | 7.97 | 88 | 3.2 | 30.3 | 989 | 440 |
| STET | Sandy loam | 70 | 12 | 18 | 88 | 5 | 30.8 | 1211 | 535 |
| Santiago | Sandy loam | 54 | 26 | 20 | 2573 | 13.3 | 9.8 | 672 | 100 |
| FP2 | Sandy clay loam | 52 | 20 | 28 | 3050 | 12.5 | 8.9 | 672 | 90 |
| FLOR1 | Sandy clay loam | 59 | 19 | 22 | 2983 | 12.6 | 8.9 | 671 | 90 |

^1^ Elevation

^2^ Mean annual temperature

^3^ Continentality

^4^ Mean annual precipitation

^5^ Winter & spring precipitation

Table S3. Ecological and plant description of sampled sites.

| **Site** | **Ecological region** | **Vegetation remarks** |
| --- | --- | --- |
| AMOS | Boreal forest | Ferns, *Rubus spp*., *Epilobium spp*., seedlings of *Salix spp*., *Betula spp*., and *Abies balsamea*. |
| STFE | Boreal forest | Ferns, *Rubus spp*., *Epilobium spp*., seedlings of *Salix spp*., *Betula spp*., and *Abies balsamea*. |
| AB | Boreal forest | *Abies balsamea*, *Picea mariana*, *P. glauca*, and *Betula papyrifera*. |
| ESSI | Cold temperate forest | Mixed *P. tremuloides* with conifers. |
| FORE | Cold temperate forest | Mixed *P. tremuloides* with conifers. |
| STET | Cold temperate forest | *Populus tremuloides*. |
| Santiago | Warm temperate forest | Pine-oak forest. *Pinus arizonica, Quercus sideroxyla,* and *Arbutus spp.* |
| FP2 | Warm temperate forest | Pine-oak forest. *Arbutus bicolor, Arbutus spp., Pinus durangensis, P. arizonica*, and *Quercus rugosa*. |
| FLOR1 | Warm temperate forest | Pine-oak forest. *Arbutus bicolor, Arbutus spp., Pinus durangensis, P. arizonica*, and *Quercus rugosa*. |

Table S4. Physicochemical properties of samples from natural stands.

| **SampleID** | **Total C [%]** | **Total N [%]** | **pH H2O** | **pH CaCl2** | **P [mg/kg]** | **CEC [cmol(+)/kg]** |
| --- | --- | --- | --- | --- | --- | --- |
| ESSI.001.S.Soil | 1.03 | 0.03 | 4.23 | 3.67 | 30.99 | 12.9 |
| ESSI.002.S.Soil | 15.3 | 0.77 | 4.36 | 3.81 | 10.07 | 21.64 |
| ESSI.003.S.Soil | 1.35 | 0.06 | 5.55 | 4.85 | 30.31 | 17.89 |
| ESSI.004.S.Soil | 2.53 | 0.13 | 4.78 | 4.14 | 499.54 | 6.69 |
| ESSI.005.S.Soil | 0.7 | 0.02 | 4.6 | 3.96 | 34.75 | 13.93 |
| AMOS.001.S.Soil | 4.54 | 0.32 | 5.07 | 4.59 | 17.59 | 23.8 |
| AMOS.002.S.Soil | 4.83 | 0.31 | 5.13 | 4.66 | 12.98 | 25.13 |
| AMOS.003.S.Soil | 6.51 | 0.38 | 5.46 | 5.03 | 43.84 | 18.38 |
| AMOS.004.S.Soil | 4.33 | 0.27 | 4.85 | 4.31 | 29.05 | 27.93 |
| AMOS.005.S.Soil | 2.27 | 0.15 | 5.08 | 4.51 | 26.88 | 23.39 |
| STFE.006.S.Soil | 6.57 | 0.37 | 5.11 | 4.51 | 6.99 | 34.38 |
| STFE.007.S.Soil | 4.82 | 0.26 | 5.22 | 4.56 | 6.35 | 32.1 |
| STFE.008.S.Soil | 2.49 | 0.16 | 5.52 | 4.84 | 6.06 | 36.95 |
| STFE.009.S.Soil | 4.72 | 0.22 | 5.27 | 4.59 | 7.23 | 32.97 |
| STFE.010.S.Soil | 2.94 | 0.14 | 5.23 | 4.49 | 7.29 | 33.01 |
| FORE.001S.Soil | 1.72 | 0.1 | 5.45 | 4.54 | 50.89 | 4.36 |
| FORE.002S.Soil | 1.63 | 0.08 | 5.29 | 4.42 | 35.27 | 4.78 |
| FORE.003S.Soil | 1.37 | 0.05 | 5.12 | 4.47 | 41.12 | 3.93 |
| FORE.004S.Soil | 1.56 | 0.06 | 5.29 | 4.43 | 26.23 | 19.75 |
| FORE.005S.Soil | 0.6 | 0.03 | 5.99 | 4.6 | 45.1 | 3.74 |
| FORE.006S.Soil | 0.36 | 0 | 5.6 | 5.01 | 54.46 | 2.65 |
| STET.001.Soil | 2.3 | 0.1 | 5.96 | 5.37 | 25.92 | 20.21 |
| STET.002.Soil | 2.7 | 0.15 | 4.64 | 4.01 | 25.1 | 16.51 |
| STET.006.Soil | 4.35 | 0.19 | 5.44 | 4.74 | 31.64 | 22.61 |
| STET.007.Soil | 3.69 | 0.15 | 5.95 | 5.37 | 21.02 | 23.05 |
| STET.008.Soil | 9.57 | 0.37 | 5.17 | 4.61 | 43.33 | 11.93 |
| Santiago.1.3.Soil | 0.58 | 0.02 | 6.48 | 5.26 | 2.9 | 20.47 |
| Santiago.2.3.Soil | 1.28 | 0.04 | 5.63 | 4.71 | 5.97 | 19.82 |
| Santiago.3.3.Soil | 0.38 | 0.01 | 5.94 | 4.72 | 5.02 | 12.18 |
| Santiago.4.3.Soil | 1.23 | 0.0375 | 5.52 | 4.68 | 12.05 | 17.79 |
| Santiago.5.3.Soil | 1.9 | 0.08 | 6.06 | 5.33 | 6.81 | 22 |
| Santiago.6.3.Soil | 0.59 | 0.01 | 6.29 | 5.34 | 2.55 | 21.17 |
| Santiago.7.3.Soil | 1.69 | 0.06 | 5.11 | 4.28 | 4.29 | 21.25 |
| Santiago.8.3.Soil | 1.21 | 0.04 | 5.22 | 4.4 | 4.99 | 13.43 |
| Santiago.9.3.Soil | 1.7 | 0.07 | 5.32 | 4.45 | 12.5 | 14.72 |
| Santiago.10.3.Soil | 1.37 | 0.04 | 5.8 | 4.9 | 6.09 | 21.36 |
| FLOR1.1.3.Soil | 2.53 | 0.13 | 5.43 | 4.94 | 24.84 | 28.81 |
| FLOR1.2.3.Soil | 3.74 | 0.18 | 5.02 | 4.63 | 33.23 | 12.9 |
| **SampleID** | **Total C [%]** | **Total N [%]** | **pH H2O** | **pH CaCl2** | **P [mg/kg]** | **CEC [cmol(+)/kg]** |
| FLOR1.3.3.Soil | 4.35 | 0.17 | 5.36 | 4.71 | 17.52 | 28.75 |
| FLOR1.4.3.Soil | 10.7 | 0.71 | 4.87 | 4.6 | 85.16 | 20.57 |
| FLOR1.5.3.Soil | 7.44 | 0.34 | 5.98 | 5.6 | 127.33 | 25.58 |
| FP2.1.3.Soil | 3.18 | 0.14 | 5.76 | 5.04 | 7.43 | 26.51 |
| FP2.5.3.Soil | 2.37 | 0.11 | 5.09 | 4.41 | 6.71 | 25.64 |
| FP2.7.3.Soil | 2.78 | 0.14 | 5.91 | 5.4 | 10.35 | 32.88 |
| FP2.8.3.Soil | 2.07 | 0.11 | 5.63 | 5.05 | 17.84 | 26.46 |
| FP2.9.3.Soil | 2.99 | 0.14 | 5.7 | 5.26 | 22.21 | 30.33 |

Table S5. Macro and micronutrients of samples from natural stands. Values are given in units of cmol(+) kg^-1^.

| **SampleID** | **K** | **Ca** | **Mg** | **Mn** | **Al** | **Fe** | **Na** |
| --- | --- | --- | --- | --- | --- | --- | --- |
| ESSI.001.S.Soil | 0.05 | 0.09 | 0.08 | 0.01 | 11.17 | 1.48 | 0.03 |
| ESSI.002.S.Soil | 0.36 | 1.13 | 0.52 | 0.03 | 16.92 | 2.56 | 0.11 |
| ESSI.003.S.Soil | 0.07 | 3.15 | 0.35 | 0.06 | 12.93 | 1.23 | 0.09 |
| ESSI.004.S.Soil | 0.12 | 1.64 | 0.16 | 0.09 | 1.99 | 2.57 | 0.13 |
| ESSI.005.S.Soil | 0.07 | 0.27 | 0.11 | 0.02 | 12.31 | 1.13 | 0.02 |
| AMOS.001.S.Soil | 0.31 | 5.78 | 1.44 | 0.34 | 14.49 | 1.37 | 0.07 |
| AMOS.002.S.Soil | 0.71 | 5.92 | 1.98 | 0.18 | 14.74 | 1.51 | 0.1 |
| AMOS.003.S.Soil | 0.86 | 10.95 | 2.67 | 0.19 | 2.01 | 1.66 | 0.03 |
| AMOS.004.S.Soil | 0.36 | 3.1 | 1.03 | 0.25 | 21.42 | 1.67 | 0.1 |
| AMOS.005.S.Soil | 0.21 | 5.31 | 2.29 | 0.17 | 13.56 | 1.76 | 0.09 |
| STFE.006.S.Soil | 0.61 | 12.68 | 3.2 | 0.06 | 15.25 | 2.46 | 0.12 |
| STFE.007.S.Soil | 0.38 | 10.96 | 3.34 | 0.03 | 15 | 2.25 | 0.14 |
| STFE.008.S.Soil | 0.63 | 11.63 | 3.2 | 0.29 | 18.98 | 2.08 | 0.14 |
| STFE.009.S.Soil | 0.65 | 11.09 | 2.95 | 0.05 | 15.59 | 2.51 | 0.13 |
| STFE.010.S.Soil | 0.33 | 8.91 | 2.56 | 0.05 | 18.45 | 2.49 | 0.23 |
| FORE.001S.Soil | 0.14 | 0.87 | 0.1 | 0.03 | 1.95 | 1.13 | 0.14 |
| FORE.002S.Soil | 0.15 | 1.01 | 0.21 | 0.02 | 1.97 | 1.26 | 0.16 |
| FORE.003S.Soil | 0.13 | 0.28 | 0.06 | 0.06 | 1.98 | 1.33 | 0.07 |
| FORE.004S.Soil | 0.11 | 0.58 | 0.08 | 0.01 | 17.24 | 1.58 | 0.14 |
| FORE.005S.Soil | 0.09 | 0.57 | 0.08 | 0.05 | 2.03 | 0.66 | 0.27 |
| FORE.006S.Soil | 0.04 | 0.15 | 0.02 | 0.02 | 2 | 0.39 | 0.02 |
| STET.001.Soil | 0.15 | 6.45 | 0.53 | 0.13 | 11.65 | 1.28 | 0.03 |
| STET.002.Soil | 0.16 | 1.31 | 0.32 | 0.07 | 13.42 | 1.22 | 0.01 |
| STET.006.Soil | 0.13 | 4.91 | 0.34 | 0.17 | 15.85 | 1.19 | 0.02 |
| STET.007.Soil | 0.14 | 7.87 | 0.46 | 0.14 | 13.4 | 1.03 | 0.02 |
| STET.008.Soil | 0.22 | 7.71 | 0.7 | 0.02 | 2 | 1.23 | 0.05 |
| Santiago.1.3.Soil | 0.35 | 11.46 | 2.06 | 0.27 | 5.94 | 0.34 | 0.04 |
| Santiago.2.3.Soil | 0.38 | 10.55 | 2.5 | 0.19 | 5.73 | 0.42 | 0.04 |
| Santiago.3.3.Soil | 0.68 | 4.61 | 0.94 | 0.49 | 5.17 | 0.24 | 0.04 |
| Santiago.4.3.Soil | 0.5 | 5.04 | 1.22 | 0.31 | 9.71 | 1 | 0.01 |
| Santiago.5.3.Soil | 0.91 | 9.99 | 2.31 | 0.88 | 7.07 | 0.82 | 0.02 |
| Santiago.6.3.Soil | 0.44 | 12.11 | 2.67 | 0.21 | 5.44 | 0.26 | 0.05 |
| Santiago.7.3.Soil | 0.68 | 6.34 | 1.68 | 0.34 | 11.14 | 1.05 | 0.03 |
| Santiago.8.3.Soil | 0.34 | 4.03 | 1.1 | 0.28 | 7.09 | 0.57 | 0.03 |
| Santiago.9.3.Soil | 0.82 | 4.95 | 1.12 | 0.4 | 6.65 | 0.76 | 0.02 |
| Santiago.10.3.Soil | 0.36 | 11.04 | 2.47 | 0.41 | 6.38 | 0.65 | 0.04 |
| FLOR1.1.3.Soil | 0.76 | 8.2 | 1.04 | 0.22 | 17.6 | 0.96 | 0.02 |
| FLOR1.2.3.Soil | 0.6 | 8.12 | 0.73 | 0.34 | 2.1 | 0.94 | 0.08 |
| **SampleID** | **K** | **Ca** | **Mg** | **Mn** | **Al** | **Fe** | **Na** |
| FLOR1.3.3.Soil | 0.47 | 8.67 | 0.88 | 0.19 | 16.16 | 2.24 | 0.13 |
| FLOR1.4.3.Soil | 1.03 | 14.68 | 1.13 | 0.7 | 2.03 | 0.94 | 0.05 |
| FLOR1.5.3.Soil | 1.68 | 18.89 | 1.59 | 0.48 | 2.03 | 0.85 | 0.07 |
| FP2.1.3.Soil | 0.94 | 8.85 | 1.45 | 0.56 | 13.67 | 0.98 | 0.07 |
| FP2.5.3.Soil | 0.76 | 4.92 | 0.97 | 0.28 | 17.67 | 1.02 | 0.02 |
| FP2.7.3.Soil | 0.56 | 12.21 | 0.99 | 0.31 | 18.02 | 0.75 | 0.03 |
| FP2.8.3.Soil | 0.93 | 8.35 | 1.5 | 0.41 | 14.31 | 0.93 | 0.02 |
| FP2.9.3.Soil | 1.09 | 12.89 | 1.53 | 0.34 | 13.23 | 1.23 | 0.02 |

Table S6. Envfit analysis measuring correlation between soil properties and ordination of bacterial/archaeal and fungal microbial structure in the complete dataset.

|  | **Bacteria/Archaea** | | | | **Fungi** | | | |
| --- | --- | --- | --- | --- | --- | --- | --- | --- |
| **Factor** | **PCo 1** | **PCo 2** | **r^2^** | ***P*-value** | **PCo 1** | **PCo 2** | **r^2^** | ***P*-value** |
| **Clay** | -0.557 | -0.830 | **0.70** | 0.001 | -0.682 | -0.732 | **0.25** | 0.010 |
| **Sand** | 0.420 | 0.908 | **0.60** | 0.001 | 0.539 | 0.842 | **0.21** | 0.017 |
| **Fe** | -1.000 | -0.026 | **0.56** | 0.001 | -0.873 | 0.488 | 0.11 | 0.084 |
| **Ca** | 0.270 | -0.963 | **0.54** | 0.001 | -0.106 | -0.994 | **0.44** | 0.001 |
| **Mg** | -0.196 | -0.981 | **0.50** | 0.001 | -0.402 | -0.916 | **0.29** | 0.003 |
| **Mn** | 0.889 | -0.458 | **0.48** | 0.001 | 0.420 | -0.907 | **0.45** | 0.001 |
| **pH** | 0.665 | -0.747 | **0.44** | 0.001 | 0.166 | -0.986 | 0.10 | 0.121 |
| **CEC** | -0.154 | -0.988 | **0.42** | 0.001 | -0.239 | -0.971 | **0.38** | 0.002 |
| **Na** | -0.969 | 0.248 | **0.35** | 0.001 | -0.428 | 0.904 | **0.16** | 0.036 |
| **K** | 0.713 | -0.702 | **0.32** | 0.003 | 0.321 | -0.947 | **0.48** | 0.001 |
| **Silt** | 0.183 | -0.983 | **0.30** | 0.001 | 0.087 | -0.996 | 0.12 | 0.089 |
| **Al** | -0.683 | -0.730 | **0.13** | 0.045 | -0.427 | -0.904 | 0.09 | 0.180 |
| **C:N ratio** | 0.716 | 0.698 | 0.12 | 0.053 | -0.466 | 0.885 | 0.00 | 0.909 |
| **total N** | -0.894 | -0.448 | 0.11 | 0.088 | -0.774 | -0.633 | **0.15** | 0.048 |
| **total C** | -0.915 | -0.403 | 0.09 | 0.135 | -0.720 | -0.694 | **0.14** | 0.045 |
| **P** | -0.197 | 0.980 | 0.08 | 0.147 | 0.053 | 0.999 | 0.06 | 0.230 |

Table S7. Envfit analysis measuring correlation between soil properties and ordination of bacterial/archaeal and fungal microbial structure in Canadian genetic group.

|  | **Bacteria/Archaea** | | | | **Fungi** | | | |
| --- | --- | --- | --- | --- | --- | --- | --- | --- |
| **Factor** | **PCo 1** | **PCo 2** | **r^2^** | ***P*-value** | **PCo 1** | **PCo 2** | **r^2^** | ***P*-value** |
| **Ca** | 0.95 | 0.31 | **0.88** | 0.001 | -0.64 | -0.77 | **0.41** | 0.004 |
| **Mg** | 0.88 | 0.47 | **0.79** | 0.001 | -0.12 | -0.99 | **0.33** | 0.016 |
| **Clay** | 0.96 | 0.28 | **0.75** | 0.001 | 0.41 | -0.91 | **0.38** | 0.007 |
| **CEC** | 0.91 | 0.40 | **0.65** | 0.001 | -0.16 | -0.99 | **0.46** | 0.004 |
| **Sand** | -0.99 | -0.14 | **0.65** | 0.001 | -0.53 | 0.85 | **0.37** | 0.008 |
| **Mn** | 0.70 | -0.71 | **0.51** | 0.003 | 0.68 | -0.73 | **0.41** | 0.004 |
| **K** | 0.98 | 0.20 | **0.44** | 0.004 | 0.43 | -0.90 | **0.25** | 0.047 |
| **Fe** | 0.52 | 0.86 | **0.39** | 0.005 | -1.00 | 0.01 | 0.13 | 0.183 |
| **Silt** | 0.92 | -0.39 | **0.35** | 0.006 | 0.68 | -0.73 | **0.33** | 0.013 |
| **Al** | 0.89 | 0.46 | 0.20 | 0.093 | 0.12 | -0.99 | **0.28** | 0.030 |
| **Na** | 0.06 | 1.00 | 0.19 | 0.075 | -0.24 | 0.97 | 0.01 | 0.921 |
| **C:N ratio** | -0.96 | 0.28 | 0.18 | 0.076 | -0.74 | 0.67 | 0.11 | 0.269 |
| **pH** | 0.83 | -0.56 | 0.16 | 0.140 | -0.90 | 0.44 | 0.03 | 0.710 |
| **total N** | 1.00 | 0.03 | 0.11 | 0.256 | 0.04 | -1.00 | 0.21 | 0.063 |
| **P** | -0.97 | -0.26 | 0.08 | 0.384 | 0.50 | 0.87 | 0.05 | 0.568 |
| **total C** | 0.97 | 0.24 | 0.07 | 0.411 | -0.92 | -0.39 | 0.20 | 0.072 |

Table S8. Envfit analysis measuring correlation between soil properties and ordination of bacterial/archaeal and fungal microbial structure in Mexican genetic group.

|  | **Bacteria/Archaea** | | | | **Fungi** | | | |
| --- | --- | --- | --- | --- | --- | --- | --- | --- |
| **Factor** | **PCo 1** | **PCo 2** | **r^2^** | ***P*-value** | **PCo 1** | **PCo 2** | **r^2^** | ***P*-value** |
| **Silt** | 0.56 | 0.83 | **0.71** | 0.002 | 0.06 | 1.00 | 0.23 | 0.168 |
| **pH** | 0.91 | -0.41 | **0.68** | 0.001 | -0.37 | -0.93 | 0.07 | 0.578 |
| **CEC** | -0.29 | -0.96 | **0.60** | 0.002 | 0.12 | -0.99 | 0.08 | 0.564 |
| **C:N ratio** | 0.89 | 0.46 | **0.59** | 0.002 | 0.08 | 1.00 | 0.26 | 0.130 |
| **Ca** | 0.36 | -0.93 | **0.53** | 0.001 | -0.30 | -0.95 | 0.22 | 0.208 |
| **Clay** | -0.67 | -0.74 | **0.51** | 0.005 | -0.10 | -0.99 | 0.24 | 0.159 |
| **Fe** | -0.91 | -0.42 | **0.47** | 0.001 | 0.51 | 0.86 | **0.36** | 0.030 |
| **K** | -0.49 | -0.87 | **0.40** | 0.009 | -0.13 | -0.99 | 0.17 | 0.257 |
| **Mg** | 1.00 | -0.05 | **0.39** | 0.012 | -0.22 | 0.98 | 0.04 | 0.742 |
| **Al** | -0.84 | -0.54 | **0.31** | 0.046 | 0.94 | 0.35 | 0.17 | 0.269 |
| **total C** | -0.67 | -0.74 | 0.23 | 0.100 | -0.23 | -0.97 | 0.09 | 0.537 |
| **P** | -0.22 | -0.98 | 0.20 | 0.136 | -0.25 | -0.97 | 0.15 | 0.338 |
| **total N** | -0.74 | -0.67 | 0.17 | 0.200 | -0.28 | -0.96 | 0.11 | 0.488 |
| **Sand** | 0.00 | -1.00 | 0.04 | 0.688 | 0.44 | 0.90 | 0.01 | 0.921 |
| **Na** | 0.30 | -0.95 | 0.04 | 0.755 | 0.21 | 0.98 | 0.03 | 0.784 |
| **Mn** | -0.73 | -0.68 | 0.00 | 0.956 | -0.27 | -0.96 | 0.11 | 0.420 |

Table S9. Coefficients, 95% confidence intervals, *P*-values and pseudo *R*^2^ for the explanatory terms retained in the selected beta regression model explaining Gini-Simpson in bacteria/Archaea.

| **Variable** | **Estimate** | **Std Error** | **z value** | ***P*-value** | **CI  2.5 %** | **CI  97.5 %** | **Pseudo *R*^2^** |
| --- | --- | --- | --- | --- | --- | --- | --- |
| Intercept | 9.49 | 3.98 | 2.38 | 0.017 | 1.69 | 17.29 |  |
| Mexico | 11.36 | 6.92 | 1.64 | 0.100 | -2.19 | 24.92 | 0.03 (group) |
| **Na** | **-3.12** | **0.92** | **-3.41** | **0.001** | **-4.92** | **-1.33** | 0.17 (Na) |
| Canada : C:N | -0.22 | 0.16 | -1.41 | 0.159 | -0.52 | 0.09 | 0.11  (group : C:N) |
| **Mexico : C:N** | **-0.63** | **0.25** | **-2.53** | **0.011** | **-1.12** | **-0.14** |  |
| Canada : pH | -0.64 | 0.80 | -0.80 | 0.425 | -2.20 | 0.93 | 0.12  (group : pH) |
| **Mexico : pH** | **-2.96** | **1.02** | **-2.89** | **0.004** | **-4.97** | **-0.96** |  |
| Canada : C:N : pH | 0.04 | 0.03 | 1.16 | 0.248 | -0.03 | 0.10 | 0.11  (group : C:N : pH) |
| **Mexico : C:N : pH** | **0.12** | **0.05** | **2.63** | **0.009** | **0.03** | **0.21** |  |
| phi | 1881.4 | 424.1 | 4.44 | 9.15E-06 | 1050.2 | 2712.6 |  |
| *Full model* |  |  |  |  |  |  | 0.58 |
